# Phylogeny-metabolism dual-directed single-cell genomics for dissecting and mining ecosystem function

**DOI:** 10.1101/2023.11.27.568714

**Authors:** Xiaoyan Jing, Yanhai Gong, Zhidian Diao, Yan Ma, Yu Meng, Jie Chen, Yishang Ren, Yinchao Li, Weihan Sun, Jia Zhang, Yuetong Ji, Yuting Liang, Zhiqi Cong, Shengying Li, Bo Ma, Zhisong Cui, Li Ma, Jian Xu

**Affiliations:** Single-Cell Center, CAS Key Laboratory of Biofuels, Qingdao Institute of BioEnergy and Bioprocess Technology, Chinese Academy of Sciences, Qingdao, China; Department of Biotechnology, College of Life Science and Technology, Huazhong University of Science and Technology, Wuhan, Hubei, China; Marine Bioresource and Environment Research Center, Key Laboratory of Marine Eco-Environmental Science and Technology, First Institute of Oceanography, Ministry of Natural Resources of China, Qingdao, Shandong, China; State Key Laboratory of Microbial Technology, Shandong University, Qingdao, Shandong, China; State Key Laboratory of Soil and Sustainable Agriculture, Institute of Soil Science, Chinese Academy of Science, Nanjing, Jiangsu, China; University of Chinese Academy of Sciences, Beijing, China; Shandong Energy Institute, Qingdao, Shandong, China; Qingdao New Energy Shandong Laboratory, Qingdao, Shandong, China; Qingdao Single-Cell Biotechnology, Co., Ltd., Qingdao, Shandong, China

**Keywords:** Fluorescence In-Situ Hybridization (FISH), Single-cell Raman microspectroscopy, Raman-activated Cell Sorting (RACS), microbiota, pollutant degradation

## Abstract

Although microbiome-wide association studies (MWAS) have uncovered many marker organisms for an ecosystem trait, mechanisms of most microbiota-mediated processes remain elusive, due to challenges in validating the markers’ *in situ* metabolic activities and tracing such activities to individual genomes. Here we introduced a phylogeny-metabolism dual-directed single-cell genomics approach called Fluorescence-In-Situ-Hybridization-guided Single-Cell Raman-activated Sorting and Sequencing (FISH-scRACS-Seq). It directly localizes individual cells from target taxon via a FISH probe for marker organism, profiles their *in situ* metabolic functions via single-cell Raman spectra, sorts cells of target taxonomy and target metabolism, and produces indexed, high-coverage and precisely-one-cell genomes. From cyclohexane-contaminated seawater, cells representing the MWAS-derived marker taxon of γ-Proteobacteria and that are actively degrading cyclohexane *in situ* were directly identified via FISH and Raman respectively, then sorted and sequenced for one-cell full genomes. In such a *Pseudoalteromonas fuliginea* cell, we discovered a three-component cytochrome P450 system that can convert cyclohexane to cyclohexanol *in vitro*, representing a previously unknown group of cyclohexane-degrading enzymes and organisms. By culture-independently unveiling enzymes, pathways, genomes and their *in situ* functions specifically for those single-cells with ecological relevance, FISH-scRACS-Seq is a rational and generally applicable approach for dissecting and mining microbiota functions.

**Teaser:** FISH-scRACS-Seq is a new strategy to dissect microbiota functional mechanism at single-cell resolution.

## Introduction

Via their rich and diverse metabolic activities, microbial consortia have supported most if not all of the critical ecological processes on Earth, such as geochemical cycling of elements, environmental remediation and nutrient utilization in hosts (*1-3*). To dissect their functioning mechanisms, and also mine the underlying bioresources (e.g., useful chassis cells or enzymes), two major strategies are usually adopted. One is driven by “genotype”: for example, microbiome-wide association studies (MWAS) identify for an ecosystem trait the DNA-sequence based taxonomical or functional-gene markers, by probing correlation between metagenomes and the trait (*4, 5*). One strength of such undirected approaches is the high-throughput and exhaustive discovery of ecosystem-trait associated organisms or genes, which ensures ecological significance of these markers. However, due to the lack of information for metabolic activities and the challenge in reconstructing individual genomes from those highly heterogeneous metagenomes, it is usually difficult to validate the *in situ* functions of marker taxa and to trace the functions to responsible genomes, pathways or enzymes. Consequentially, although numerous taxonomic markers were reported for many microbiota-mediated processes, their functioning mechanisms remain elusive (*6, 7*), particularly for those involving not-yet-cultured marker organisms.

The other strategy starts with “metabolic phenotype”: for example, Raman-activated Cell Sorting and Sequencing (RACS-Seq), a metabolism-directed approach, can directly identify individual cells of target metabolism that corresponds to the ecosystem trait, and then track the metabolic activity to the underpinning single-cell genomes (*8-11*). Specifically, individual cells of microbiota are profiled for single-cell Raman spectra (SCRS) which serve as a proxy of *in situ* metabolic phenome (*11*), and those cells of target phenotypes are sorted via a RACS instrument and then sequenced for their single-cell full genomes at an indexed, one-cell-one-tube manner (*11-13*). When coupled with stable isotope probing (SIP-Raman), SCRS can measure cellular intake rate of substrates (e.g., ^13^C, ^15^N, ^18^O and D (*8, 13-19*)), and the D_2_O-intake based cellular vitality can be used to model degradative activity of carbon source (*16, 20, 21*). In addition, SCRS can reveal the biosynthetic profile of cell (e.g., carotenoids (*22-25*), proteins (*26*), triacylglycerols (*26-28*) and other Raman-sensitive compounds), and characterize cellular response to environmental changes (e.g., susceptibility to drugs or other types of stresses (*11, 29, 30*)). A core strength of this strategy is the ability to actually measure *in situ* metabolic activity and directly trace it to genomes, both at single-cell resolution (*24, 25, 29, 31-33*). However, due to the sheer number of cells in microbiota yet the comparably low throughput of RACS-Seq at present, such metabolism-directed single-cell genomics approach is usually shallow in sampling depth and narrow in investigative scope. As a result, *in situ* metabolism of those selected cells of interest such as marker organisms identified by MWAS, cannot be probed in a targeted manner, i.e., whether the single-cell metabolic phenomes and genomes produced by RACS-Seq are of ecological relevance is usually not clear.

To tackle these challenges, we introduce a phylogeny-metabolism dual-directed single-cell omics approach called Fluorescence-In-Situ-Hybridization-guided Single-Cell Raman-activated Sorting and Sequencing (FISH-scRACS-Seq). Based on a FISH probe designed from MWAS-derived taxonomical markers, the method directly localizes individual cells of the target taxon in a microbiota, profiles their *in situ* metabolic functions via SCRS, then sorts for those cells of target taxonomy and target metabolism, and finally produces their indexed, high-coverage and precisely-one-bacterial-cell genomes. The method was evaluated via a series of mock community and then demonstrated for soil and seawater microbiota. Coupling of FISH-scRACS-Seq to upstream MWAS allowed efficient, culture-independent tracing of cyclohexane degradation in cycloalkane-contaminated seawater from a condensate gas field to the one-cell genomes of uncultured *Pseudoalteromonas fuliginea* and further to a previously unknown group of cytochrome P450 based cyclohexane monooxygenases that can convert cyclohexane to cyclohexanol *in vitro*. Therefore, FISH-scRACS-Seq is a rational and generally applicable strategy for dissecting and mining microbiota function.

## Results

### Overview of FISH-scRACS-Seq for phylogeny-metabolism dual-directed single-cell omics

The FISH-scRACS-Seq workflow for dissecting a microbiota consists of three steps (**Fig. 1**). In Step 1 (i.e., “FISH”), individual cells of a target taxon are directly localized in a microbiota sample, via a taxon-specific catalyzed reporter deposition (CARD)-FISH probe (the use of CARD-FISH greatly increase signal-noise ratio (*34-36*)). Importantly, the FISH process usually sacrifices cellular vitality (*37, 38*), therefore, prior to probe hybridization, pretreatments that support downstream SCRS-based profiling of metabolic phenome should ensue, such as the feeding of stable-isotope labelled substrates (e.g., D_2_O, H^13^CO_3_^-^, ^15^N_2_, etc (*11, 39*)).

**Figure 1.**
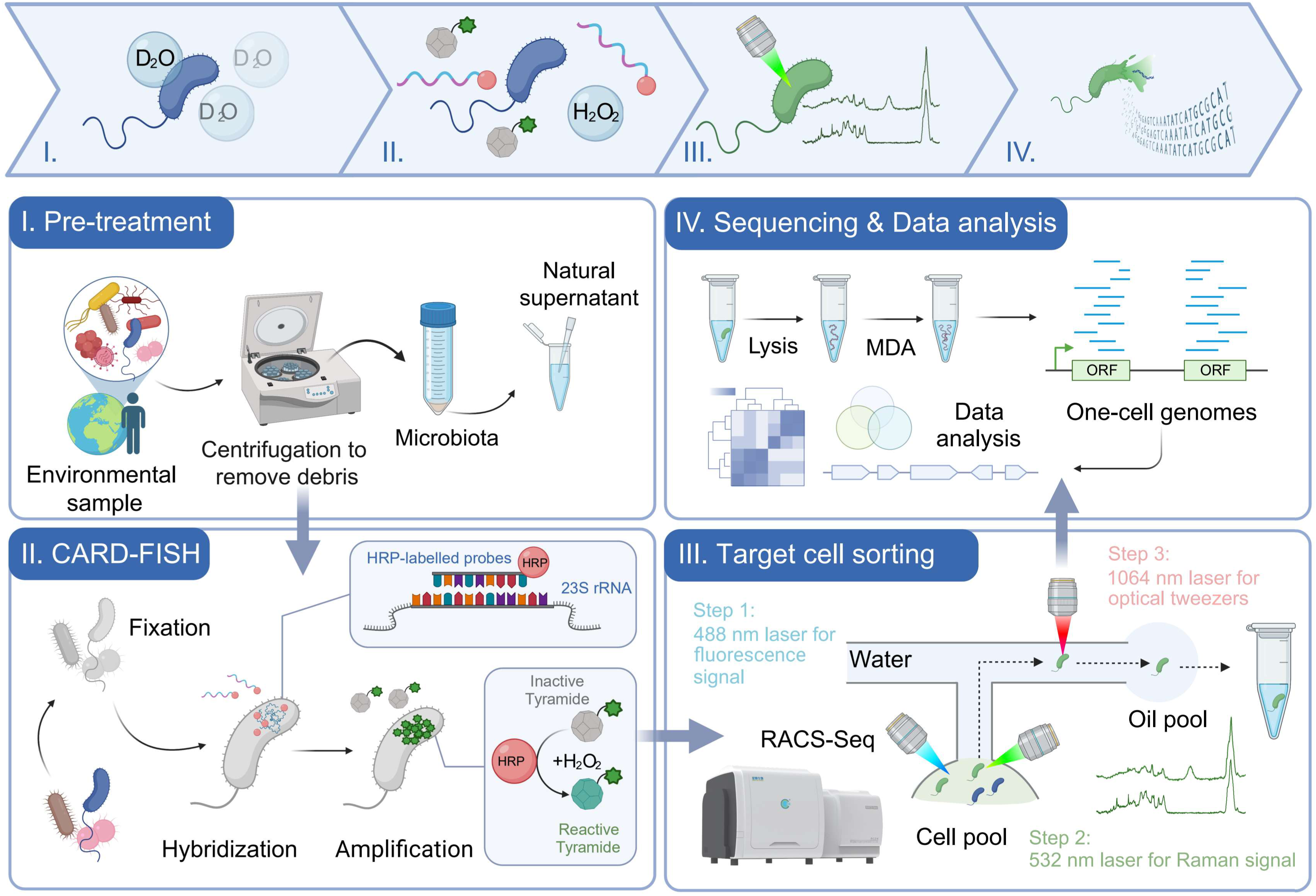
The FISH-scRACS-Seq strategy for taxonomy-guided, in-situ-function-driven profiling of microbiota at single-cell resolution. **I.** Pre-treatment: microbial extracts from environment, and then the microbial cells were labelled by D_2_O. **II.** CARD-FISH: CARD-FISH labelling was performed by fixation, hybridization and amplification successively via the taxon-specific CARD-FISH probe for γ-Proteobacteria. **III.** Raman-activated sorting of FISH-labelled cells: Fluorescence signal recognition for γ-Proteobacteria was performed via 488 nm laser; their corresponding metabolic vitality revealed via D_2_O-labeling was recognized via Raman microspectroscopy; the target cells were then sorted out in the RAGE chip, as a one-cell-encapsulated droplet. **IV.** Sequencing & data analysis: The sorted cells were lysed and its genomic DNA were amplified by MDA, then the MDA products underwent single-cell genome sequencing and analysis.

In Step 2 (i.e., “scRACS”), post-FISH cells are distinguished and sorted based on not just the target phylogeny (via the FISH probe) but also the target metabolic phenome (via the SCRS). Specifically, under aqueous suspension, the CARD-FISH-labeled cells are trapped and analyzed for SCRS individually in a RAGE chip via a 532 nm laser, which generates high signal-to-noise ratio SCRS. Then, via a technique called single-cell Raman-activated Gravity-driven Encapsulation coupled with Sequencing (scRAGE-Seq) in a RACS-Seq instrument (*31*) (**Methods**), those CARD-FISH-labeled cells with characteristic SCRS that correspond to target metabolic phenotypes are individually captured and moved with a 1,064 nm laser to form one-cell-encapsulated droplets that are then sequentially exported.

In Step 3 (i.e., “Seq”), the post-FISH-RACS cells in droplets would undergo cell lysis, Multiple Displacement Amplification (MDA), and genome sequencing in an indexed, one-cell-one-tube manner (**Fig. 1**). Notably, as the one-cell-droplets already carry an oil phase (mineral oil), an emulsion reaction for MDA would be formed simply via vortexing the tube after introducing the lysis buffer. After quality assessment, the one-cell MDA products are then shotgun sequenced individually, followed by *de novo* assembly and *in silico* genome analysis. In this way, specifically for those cells of target phylogeny in a microbiota, the target metabolic activity *in situ* is directly traced to genome sequence at single-cell resolution.

### Validation of FISH-scRACS-Seq using pure-cultured *Escherichia coli* cells

To benchmark method performance, we started from a pure culture of *E. coli* K-12 DH5α. To simulate SCRS-based profiling of metabolic phenome from a microbiota, prior to CARD-FISH labeling, the cells were first fed 50% D_2_O (vol/vol), producing a broad Raman band that peaks at 2,157 cm^-1^ in the region between 2,040 and 2,300 cm^-1^. Representing the C-D stretching vibrations shifted from the C-H stretching vibrations at 2800-3200 cm^-1^ (*16*), this C-D band can quantitatively model metabolic vitality of the cell (*15, 16*). Then the CARD-FISH process ensued, where all the *E. coli* cells were successfully labeled with the GAM42a probe, a fluorescence probe targeting γ*-*Proteobacteria (**Fig. 2A**). Then based on the specific C-D bands in SCRS (**Fig. 2B**) and the taxon-specific fluorescence signal, cells were sorted via scRAGE-Seq in a one-cell-per-tube manner (one cell-free reaction as negative control in each batch of experiments). To validate successful MDA for each sorted cell, the 16S rRNA gene was PCR-amplified using the MDA product as template (**Table S1**; **Fig. S1A**). From totally 20 target cells, twelve one-cell MDA products with both prominent MDA bands and positive 16S rRNA gene PCR results were obtained for 16S-amplicon (Sanger) and whole-genome shotgun sequencing (**WGS**; **Fig. S1A**). For the one-cell WGS, ∼1 Gb of raw sequencing data was produced for each cell (E04, E07, E08, E09, E10, E11, E12, E13, E14 and E16; the other two failed to yield sequencing libraries due to severe degradation; **Table S2**).

**Figure 2.**
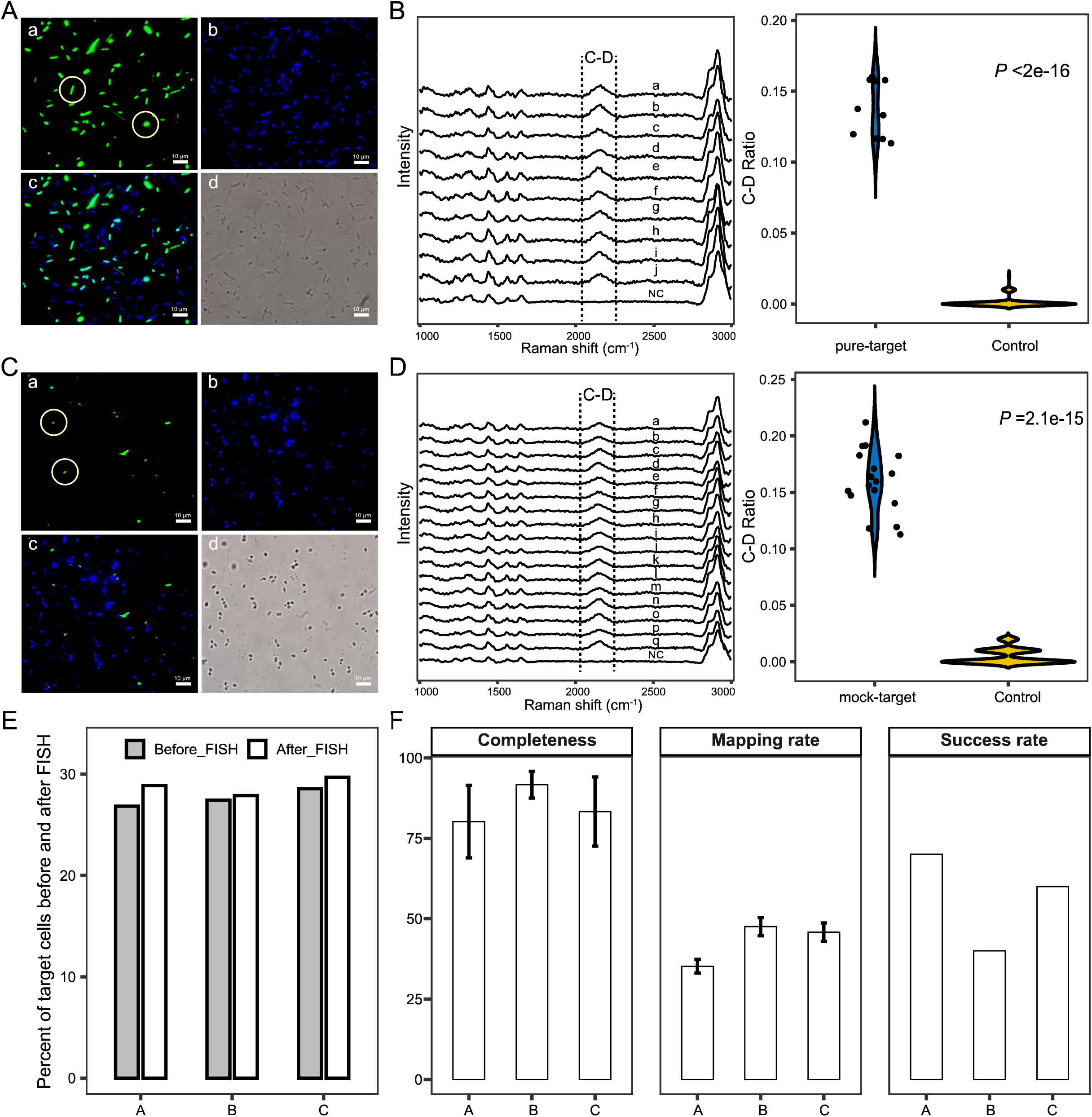
FISH-scRACS-Seq accurately and efficiently recovers SAGs from metabolically active, pure-cultured γ-Proteobacteria cells. (**A**) Photomicrographs of CARD-FISH-stained pure-cultured *Escherichia coli* K-12 DH5α. (a) Photomicrographs of cells hybridized with HRP-labelled oligonucleotide probes GAM42a (green); (b) DAPI staining (blue) of cells shown via a color-combined image recorded by epifluorescence microscopy (c); (d) phase-contrast photomicrograph. Panel (b) is the negative control (i.e., without any probes). Scale bar, 10 μm. (**B**) The C-D peaks (left panel) and their corresponding C-D Ratios (right panel) of the target cells which were sorted via both “taxon-specific” and “metabolic” phenotypes of *Escherichia coli* K-12 DH5α for single-cell genomes. Letters “a-j” represent cells with C-D peaks in SCRSs of samples “E04, E07, E08, E09, E10, E11, E12, E13, E14, and E16”, respectively, whereas the “control” SCRSs represent cells without C-D bands. (**C**) CARD-FISH Photomicrographs of four-species mock microbiota hybridized with probe GAM42a. Each series shows identical microscopic fields. Panels a to d show photomicrograph of the mock bacteria hybridized with γ-Proteobacteria targeting probe GAM42a, DAPI staining of DNA, overlay images of probe signal (green) and DAPI staining (blue), phase-contrast image, respectively. Scale bar: 10 μm. (**D**) Statistics on the percentage of the target *Escherichia coli* K-12 DH5α cells in this four-species mock microbiota before and after the CARD-FISH experiment. Three batches of experiments (A, B and C) were performed. (**E**) The C-D peaks (left panel) and their corresponding C-D Ratios (right panel) of the target cells from these four species, which were mixed for cell sorting via both “taxon-specific” and “metabolism-specific” features for single-cell genomes. (**F**) Performance validation of FISH-scRACS-Seq via statistical analysis of FISH-scRACS-derived SAGs. Three batches of experiments (A, B and C) were performed. Completeness is shown as the percentage of bases with sequencing reads and total bases in the reference genome. Mapping rate is shown as the percentage of sequencing reads which can be mapped to the reference genome. Success rate is shown as “the number of successful runs/total number of attempted runs”. Success was defined based on sequence-based verification of 16S rDNA genes amplified from the gene-specific primer pairs.

Mapping of the WGS reads to *E. coli* K-12 reference genome revealed 55.42% to 87.80% one-cell genome coverage, confirming accurate sorting and sequencing for the one-cell-per-tube reactions. Moreover, completeness of one-cell genome assemblies ranges from 65.74% to 95.53% (**Table S2**), with 60% of the SAGs from FISH-scRAGE-Seq yielding high-quality draft genomes (estimated completeness >80%; **Table S2**). Therefore, producing high-quality SCRS plus corresponding high-coverage one-cell genome from a FISH sample is feasible via FISH-scRAGE-Seq.

Another technique to sort and sequence individual cells in an indexed manner from microbiota is Raman-activated Cell Ejection (RACE-Seq), which isolates the cell of target Raman signal by ejection (via a second, 532 nm pulse laser) on a dry slide via laser induced forward transfer (*40, 41*). To compare the performance of RAGE and RACE, the ten one-cell SAGs from FISH-scRAGE-Seq were aligned to RACE-Seq derived SAGs which consists of three five-cells-pooled SAGs (one-cell RACE-Seq are not available due to the very low success-rate of scRACE-Seq; SRA accessions SRR10549451-SRR10549453 (*40*)). SAGs of FISH-scRAGE-Seq are of much higher quality than RACE-Seq in all assembly metrics (genome completeness, sequencing bias, assembly contiguity; **Fig. 3A-C**), and recovered far more genes (**Fig. 3D**, left panel). These are consistent with previous reports on low genome coverage (rarely exceeding 10% (*23, 41*)) of RACE-Seq due to premature cell vitality loss prior to Raman exposure, direct cell exposure to the Raman-exciting laser, and the influence of pulsed laser-induced cell ejection (*40*). Therefore, by operating all the FISH, Raman acquisition and sorting steps in an aquatic phase, RAGE was chosen for FISH-scRACS-Seq in subsequent experiments.

**Figure 3.**
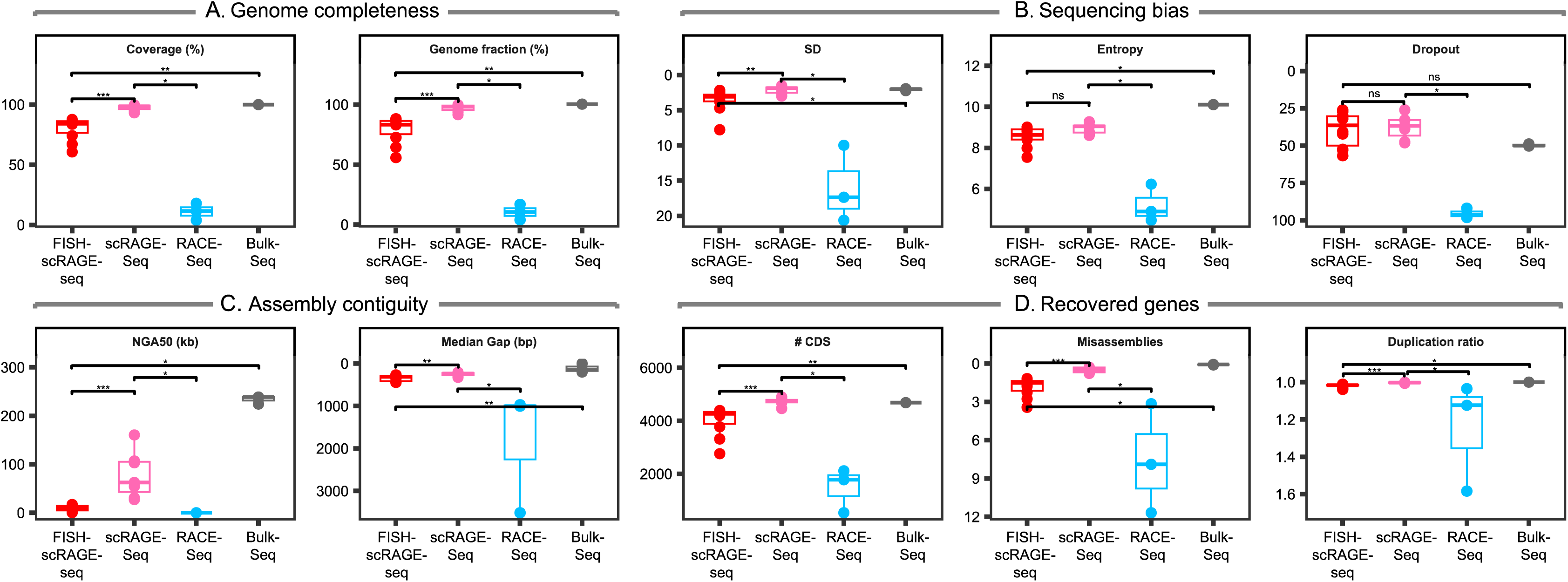
Performance of FISH-scRACS-Seq in precisely one-bacterial-cell genome sequencing. For FISH-scRACS-Seq, ten *Escherichia coli* K-12 DH5α cells (E04, E07, E08, E09, E10, E11, E12, E13, E14 and E16) were individually sorted and sequenced from precisely one cell, i.e., one SAG per cell. For scRAGE-Seq, seven *E. coli* ATCC35218 cells (X1, X2, X3, X4, X6, X7, X9) were individually sorted and sequenced, i.e., one SAG per cell. For scRACE-Seq, individual *E. coli* ATCC35218 cells were ejected by RACE separately, which generated three five-cell-pools (S1, S2, S9) for sequencing (as one-cell sorting and sequencing with RACE has suffered from low success rates), i.e., one MDA reaction per five-cell pool. In addition, three replicates of isogenic liquid culture derived from a single *E. coli* ATCC35218 colony on plate (Bulk; Ec1, Ec2, Ec3) were sequenced (each including approx. 10^9^ cells). Sequencing data from scRAGE-Seq and RACE-Seq were retrieved from NCBI SRA database (bioproject PRJNA574296 and PRJNA592282). The *de novo* assembly of Ec1 was employed as reference genome to assess the quality of aforementioned SAGs. (**A**) Completeness of the SAGs based on covered bases (left) and mapped contigs (right). (**B**) Bias in genome coverage (by sequencing reads), assessed via three parameters: SD (standard deviation of relative coverage; left), entropy (information entropy of relative coverage; middle) and drop-out rate (fraction of 200-bp fragments with relative coverage < 0.1; right). (**C**) Continuity of *de novo* assembly based on NGA50 (the length for which the collection of all aligned contigs of that length or longer covers at least half the reference genome; left) and median gap size (right). (**D**) The number of protein-coding genes mined from *de novo* assembly (left) and fidelity of *de novo* assembly, as evaluated by misassemblies (normalized by aligned contigs, middle) and duplication ratio (right).

On the other hand, scRAGE-Seq can reach one-cell whole-genome coverage of up to 100%, for microbiomes from human urine (*31*), soil (*24*), seawater (*25*), wastewater (*32*), human gastric biopsy (*29*) and probiotic products (*33*). To test whether and to what degree addition of the FISH step affects downstream scRAGE-Seq results, the ten one-cell SAGs from FISH-scRAGE-Seq were compared to scRAGE-Seq derived SAGs (seven one-cell SAGs; PRJNA574296 in the NCBI SRA database) from the same *E. coli* culture. (*i*) For read coverage, FISH-scRAGE-Seq reaches averagely ∼80.01%, which is lower than scRAGE-Seq (∼97.48%; Wilcoxon test, *p* < 0.001) but much higher than RACE-Seq (∼11.09%; **Fig. 3A**, left panel). Results from reconstructed draft genomes are similar (**Fig. 3A**, right panel). (*ii*) As for uniformity of sequence coverage, which is a major challenge in single-cell genome amplification and sequencing of microbiota (*42*), FISH-scRAGE-Seq shows similar mapping uniformity on the genomes to scRAGE-Seq and bulk, respectively (**Fig. 3B**, left panel; SD; Wilcoxon test, 3. 65 vs 2.13, *p* < 0.05; 3.65 vs 2.06, *p* < 0.01), information entropy (**Fig. 3B**, middle panel; Entroy; Wilcoxon test, 8.54 vs 8.94, *p* > 0.05; 8.54 vs 10.11, *p* < 0.05) and dropout rate (**Fig. 3B**, right panel; Dropout; Wilcoxon test, 39.20% vs 37.60%, *p* > 0.05; 39.20% vs 49.7%, *p* > 0.05). (*iii*) As for assembly continuity, FISH-scRAGE-Seq obtains shorter contigs (NGA50, 9.72 vs 78.01; Wilcoxon test, *p* < 0.001) than scRAGE-Seq (**Fig. 3C**, left panel). (*iv*) As for number of recovered genes, FISH-scRAGE-Seq recovers 86.17% genes, although 12.98% fewer than scRAGE-Seq (from the number of CDSs, 3,998 vs 4,725; Wilcoxon test, *p* < 0.001) (**Fig. 3D**, left panel), which is consistent with the lower completeness. Therefore, although introducing the FISH step can affect quality of downstream single-cell genomes, FISH-scRAGE-Seq can still produce functionally informed, high-coverage one-cell genomes for *E. coli*.

### FISH-scRACS-Seq shows high specificity and high sensitivity in dissecting a mock microbiota

To evaluate performance of FISH-scRACS-Seq (i.e., FISH-scRAGE-Seq in this case) for microbiota, we constructed a four-species mock consortium that consists of *Bacillus subtilis* H6 (*Bs*, non γ-Proteobacteria), *Escherichia coli* K-12 DH5α (*Ec*, γ-Proteobacteria), *Micrococcus luteus* D11 (*Ml*, non γ-Proteobacteria) and *Saccharomyces cerevisia*e BY4742 (*Sc*, non γ-Proteobacteria) in a 1:1:1:1 ratio (**Methods**). Sensitivity and specificity of FISH-scRACS-Seq were then evaluated by sorting the mock microbiota based on fluorescence signal (i.e., the targeted taxon) and Raman signal (i.e., the targeted metabolic function of vitality via C-D band).

We started from the FISH step, by subjecting each culture respectively to hybridization with the CARD-FISH probe of GAM42a, which specifically targets *Ec*. Microscopic examination showed that all cells in the *Ec* culture, but no cells in the other three cultures, were labelled with fluorescence (**Fig. S2**). Moreover, the four cultures of *Bs*, *Ec*, *Ml* and *Sc* were labelled with 50% D_2_O respectively prior to CARD-FISH labeling, and then mixed in a 1:1:1:1 ratio (**Fig. 2C**). Cell counting under a fluorescence microscope revealed that mean percentage of *Ec* cells before and after the FISH labelling was ∼28.8% and ∼27.6%, respectively, showing no significant difference (Wilcoxon test, *p* = 0.37; **Fig. 2D**). This is consistent with the pre-determined ratio of *Ec* in microbiota, and supports high sensitivity and specificity of CARD-FISH.

Furthermore, triplicate experiments were performed for sorting the mock microbiota not just based on fluorescence (**Fig. 2E**) but the D_2_O peak of SCRS (**Fig. 2E**; **Table 1**). In each of the three FISH-scRACS-Seq runs, ten target cells were sorted via the presence of taxon-specific fluorescence and C-D peaks, respectively. One-cell genome sequencing results suggested that (**Table 1**; **Fig. 2F**): (*i*) Sanger sequencing of all the 16S PCR products (from 17 MDA positive cells in 30 sorted cells) yield only *E. coli* specific 16S rDNA sequences, and the genome completeness of the SAGs ranges between 19.12% and 99.76%, with average of 80.18%, 91.66% and 83.30% for each run, respectively; (*ii*) 26.26% to 54.86% of the shotgun reads are mapped to *Ec*, and the average mapping rate is 35.21%, 47.53% and 45.82%, respectively; (*iii*) for each run, the success rate for a given species ranges from 40% to 70% (average of 56.67%). Notably, empty droplets derived from the aqueous phase around the target cells, which served as the negative controls, produce only negative results in 16S rRNA gene validation (**Fig. S1B-D**); these support the stringency of FISH-scRACS-Seq workflow (with the aqueous sorting microenvironment of being free of contaminating DNA from air, surface or reagents, which are often encountered in single-cell isolating and sequencing; (*43, 44*)). Collectively, results from mock microbiota also support high specificity and high sensitivity of FISH-scRACS-Seq.

**Table 1.** Benchmarking the performance of FISH-scRACS-Seq using a four-species mock microbiota, consisting of *Escherichia coli* K-12 DH5α (*Ec*), *Micrococcus luteus* D11 (*Ml*), *Bacillus subtilis* H6 (*Bs*) and *Saccharomyces cerevisiae* BY4742 (*Sc*) in a 1:1:1:1 ratio. *Ec* and *Ml* are γ**-**Proteobacteria.

| Experiment series | Sample ID | WGS reads mapped to reference genome |  |  | One-cell WGS assembly |  |  |  |  |
| --- | --- | --- | --- | --- | --- | --- | --- | --- | --- |
|  |  | Taxonomy of bins based on sequence | Mapping rate of the WGS reads (%) | Average mapping rate (%) | Genome completeness (%) | Average genome completeness (%) | Contamination (%) | WGS consistent with sorting criteria | Success rate (%) |
| **A** | **FME01** | *Ec* | 43.0 | 35.2 | 99.03 | 80.18 | 0.16 | yes | 70 |
|  | **FME03** | *Ec* | 32.7 |  | 99.38 |  | 0.25 | yes |  |
|  | **FME04** | *Ec* | 31.7 |  | 96.67 |  | 0.77 | yes |  |
|  | **FME06** | *Ec* | 34.2 |  | 83.31 |  | 5.7 | yes |  |
|  | **FME08** | *Ec* | 39.5 |  | 99.57 |  | 0.52 | yes |  |
|  | **FME09** | *Ec* | 26.3 |  | 19.12 |  | 0 | yes |  |
|  | **FME10** | *Ec* | 38.9 |  | 64.21 |  | 4.15 | yes |  |
|  | **NC_A** | - | - | - | - | - | - | - | - |
| **B** | **SME01** | *Ec* | 43.0 | 46.6 | 99.1 | 91.66 | 0.75 | yes | 40 |
|  | **SME02** | *Ec* | 39.3 |  | 80.86 |  | 2.66 | yes |  |
|  | **SME07** | *Ec* | 49.3 |  | 89.46 |  | 2.74 | yes |  |
|  | **SME08** | *Ec* | 54.7 |  | 97.21 |  | 0.84 | yes |  |
|  | **NC_B** | - | - | - | - | - | - | - | - |
| **C** | **TME02** | *Ec* | 54.8 | 45.8 | 99.76 | 83.30 | 0.78 | yes | 60 |
|  | **TME03** | *Ec* | 43.5 |  | 99.6 |  | 1.5 | yes |  |
|  | **TME05** | *Ec* | 48.3 |  | 99.35 |  | 0.21 | yes |  |
|  | **TME06** | *Ec* | 35.0 |  | 37.07 |  | 1.72 | yes |  |
|  | **TME08** | *Ec* | 50.5 |  | 98.19 |  | 0.65 | yes |  |
|  | **TME09** | *Ec* | 42.6 |  | 65.81 |  | 5.09 | yes |  |
|  | **NC_C** | - | - | - | - | - | - | - | - |

### Tests of FISH-scRACS-Seq on natural soil microbiota revealed a metabolically active *Moraxella osloensis* cell that carries a plasmid harboring antimicrobial resistance genes

To assess FISH-scRAGE-Seq’s performance in an actual environmental sample, we employed soil which harbors arguably the most metabolically and genetically heterogeneous microbiota on Earth (*45*). Samples of shallow soil were collected from grassland at a depth < 3 cm in the campus of Qingdao Institute of Bioenergy and Bioprocess Technology, Chinese Academy of Sciences, China (36°9ʹ19ʹʹN, 120°28ʹ50ʹʹE). Then, γ-Proteobacteria, which are metabolically active but of low abundance in soil and can serve as responder/indictor of soil pollution (*46-49*), were labeled via CARD-FISH as above (**Fig. 4A**). Finally, individual cells that carry the fluorescent CARD-FISH signal (i.e., of the target taxon) and the C-D peak (i.e., with the target metabolic vitality) were isolated from soil samples and sequenced via FISH-scRACS-Seq (**Fig. 4B**; **Fig. 1**; **Methods**).

**Figure 4.**
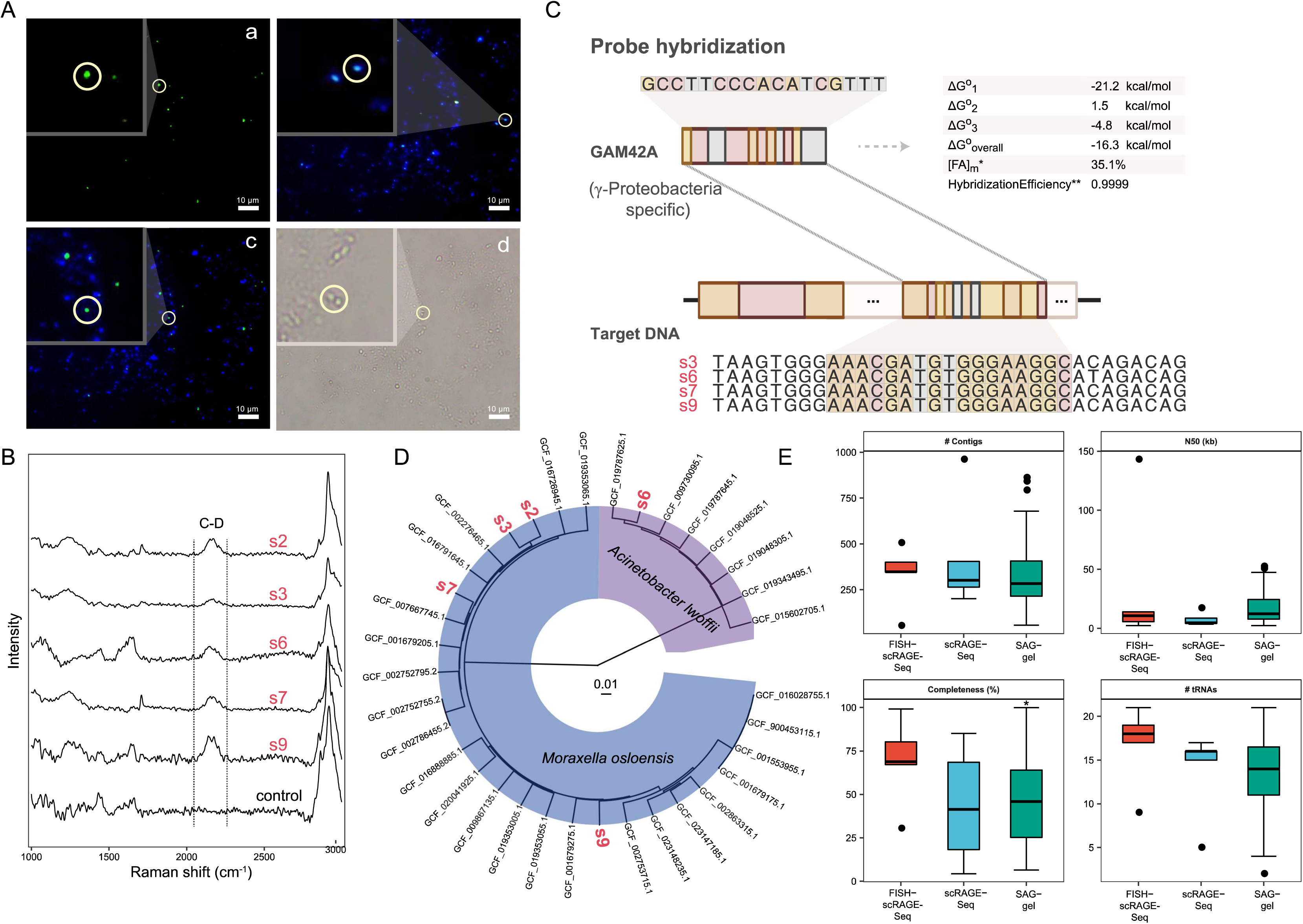
Application of FISH-scRACS-Seq in identifying, sorting and sequencing of metabolic active γ-Proteobacteria cells at one-cell resolution in soil microbiota. (**A**) Photomicrographs of γ-Proteobacteria targeted by CARD-FISH probe from soil bacteria. Panels a to d are photomicrographs of soil bacteria hybridized with γ-Proteobacteria targeting probe GAM42a, DAPI staining of DNA, overlay images of probe signal (green) and DAPI staining (blue), and phase-contrast image, respectively. Each series shows identical microscopic fields. Scale bar, 10 μm. (**B**) SCRS of the target cells in soil sample, which sorted via both “taxon-specific” and “metabolic” phenotypes of microbes for single-cell genomes. (**C**) The FISH probe perfectly matches those contained genomic regions recovered from SAGs. (**D**) Phylogenetic tree constructed using UPGMA based on ANI matrix of SAGs and collected genomes from NCBI RefSeq database. (**E**) Continuity and completeness comparison of soil-derived γ-Proteobacteria SAGs for FISH-scRAGE-Seq, scRAGE-Seq and SAG-gel. No significant differences were observed for FISH-scRAGE-Seq and scRAGE-Seq. FISH-scRAGE-Seq and SAG-gel had comparable performance, except that the completeness of FISH-Seq-derived SAGs was significantly higher than SAG-gel-derived SAGs (*p* < 0.05, Wilcoxon test).

After quality control, clean reads from the five FISH-scRAGE-Seq reactions were *de novo* assembled into five SAGs (s2, s3, s6, s7 and s9; **Table S3**). Four of the five SAGs recovered 23S rRNA gene fragments that harbor the exact hybrid site of GAM42A probe (with est. 99.99% hybridization efficiency; **Fig. 4C**). Taxonomic annotation of the five SAGs also pinpoints them as from γ-Proteobacteria (*Moraxella* spp. for s2, s3, s7, s9 and *Acinetobacter* spp. for s6; **Fig. 4D** and **Table 2**). GC% of the assembled contigs (> 200 bp; after decontamination; **Methods**) exhibit normal distribution (**Fig. S3A**). Completeness of reconstructed one-cell genomes ranges from 41.79% to 99.14% (average of ∼74.93%; **Table 2**), as estimated via lineage-specific marker genes by CheckM (*50*). These results support the feasibility of FISH-scRACS-Seq on soil microbiota.

**Table 2.** Performance of FISH-scRACS-Seq in profiling metabolic phenome and genome of γ-Proteobacteria in soil and seawater samples.

| **FISH-scRACS-sorted samples** |  | **Taxonomic classification** | **Genome completeness (%)** | **Contamination (%)** |
| --- | --- | --- | --- | --- |
| C-D peak-containing γ-Proteobacteria cells<br><u>in soil</u> | s2 | *Moraxella* | 41.79 | 7.85 |
|  | s3 | *Moraxella* | 75.60 | 6.19 |
|  | s6 | *Acinetobacter* | 85.13 | 1.57 |
|  | s7 | *Moraxella* | 99.14 | 0.10 |
|  | s9 | *Moraxella* | 72.97 | 2.38 |
| C-D peak-containing and cycloalkane-degrading γ-Proteobacteria cells <u>in seawater</u> | m1 | *Pseudoalteromonas* | 94.35 | 1.02 |
|  | m4 | *Pseudoalteromonas* | 40.49 | 2.98 |
|  | m7 | *Pseudoalteromonas* | 70.88 | 4.95 |

The FISH-scRAGE-Seq derived SAGs were further compared to scRAGE-Seq-derived (*24*) and SAG-gel-derived (*51*) γ-Proteobacteria SAGs (contigs over 1000 bp were selected for comparison; **Methods**; **Fig. 4E**). The contiguity (quantity and N50 of contigs) and completeness (genome completeness and number of unique tRNAs recovered) of SAGs among the three groups showed no significant difference (except for the completeness of FISH-scRAGE-Seq vs SAG-gel; **Fig. 4E**), thus incorporating FISH into scRACS-Seq does not degrade the quality of SAGs when analyzing complex microbiomes. Besides, in the s9 cell, the taxon-specific SAG obtained via FISH-scRAGE-Seq unraveled a plasmid that harbors a gene encoding Class A β-lactamases (**Fig. S3C**), enzymes that can inactivate β-lactam antibiotics including carbapenems. Such genes represent a major challenge in treating bacterial infections as they are highly diverse, rapidly evolving to acquire new resistance mechanisms, and easily transferred between bacteria through the spreading of plasmids (*52, 53*). Notably, this plasmid is similar in sequence (base identity of 99.5% over totally 33.0 Kb) to the plasmid 1 in *M. osloensis* strain NP7 (*54*), a γ-Proteobacteria that resides on the human skin. The sharing of such resistance-gene harboring plasmids between metabolically active γ-Proteobacteria residents from soil and those from human skin suggests *M. osloensis* as one likely reservoir and conveyor of antimicrobial resistance across the two ecosystems. Therefore, FISH-scRAGE-Seq is able to profile, directly from complex natural microbiota and in a phylogenetically directed manner, both metabolic activities and high-coverage (> 99%) genomes at precisely one-cell resolution.

### Unraveling *in situ* cycloalkane-degrading γ-Proteobacteria and their genomes at single-cell resolution in contaminated seawater by FISH-scRACS-Seq

Cyclic alkanes, abundant in both subsurface hydrocarbon reservoirs and gas condensates (*55*), are highly toxic to aquatic organisms and recalcitrant to degradation, thus pose significant ecological risks when large-scale oil spills occur (*56, 57*). Although little is known about cycloalkane biodegradation mechanism in marine ecosystems (*58*), MWAS has associated a group of uncultured, psychrophilic and oligotrophic γ-Proteobacteria to cycloalkane degradation in China’s marginal seas (*55*).

To further probe the mechanistic link between those γ-Proteobacteria and cycloalkane degradation, seven metagenomes were profiled by WGS from each of seven seawater samples of different dilution ratios. The seawater was sampled from a large condensate gas field in the Bohai Sea where cycloalkanes are released from gas mining (**Fig. 5A**). Metagenomic assembly and binning reconstructed seven MAGs of γ-Proteobacteria (phylogenetically annotated as *P. fuliginea*; one MAG in each sample; **Methods**), which are of 4.69 to 4.82 Mb in size, 99.66-100.00% in est. completeness (*42*), and contain 4,111-4,266 genes respectively. However, none of the seven MAGs encode cycloalkane degradation pathways. In fact, like many MWAS studies, due to the difficulty in measuring target taxon’s metabolic activities *in situ* and unambiguously assigning them to individual genomes, the specific organisms, pathways or enzymes responsible for the ecological trait (e.g., cycloalkane degradation) have remained speculative.

**Figure 5.**
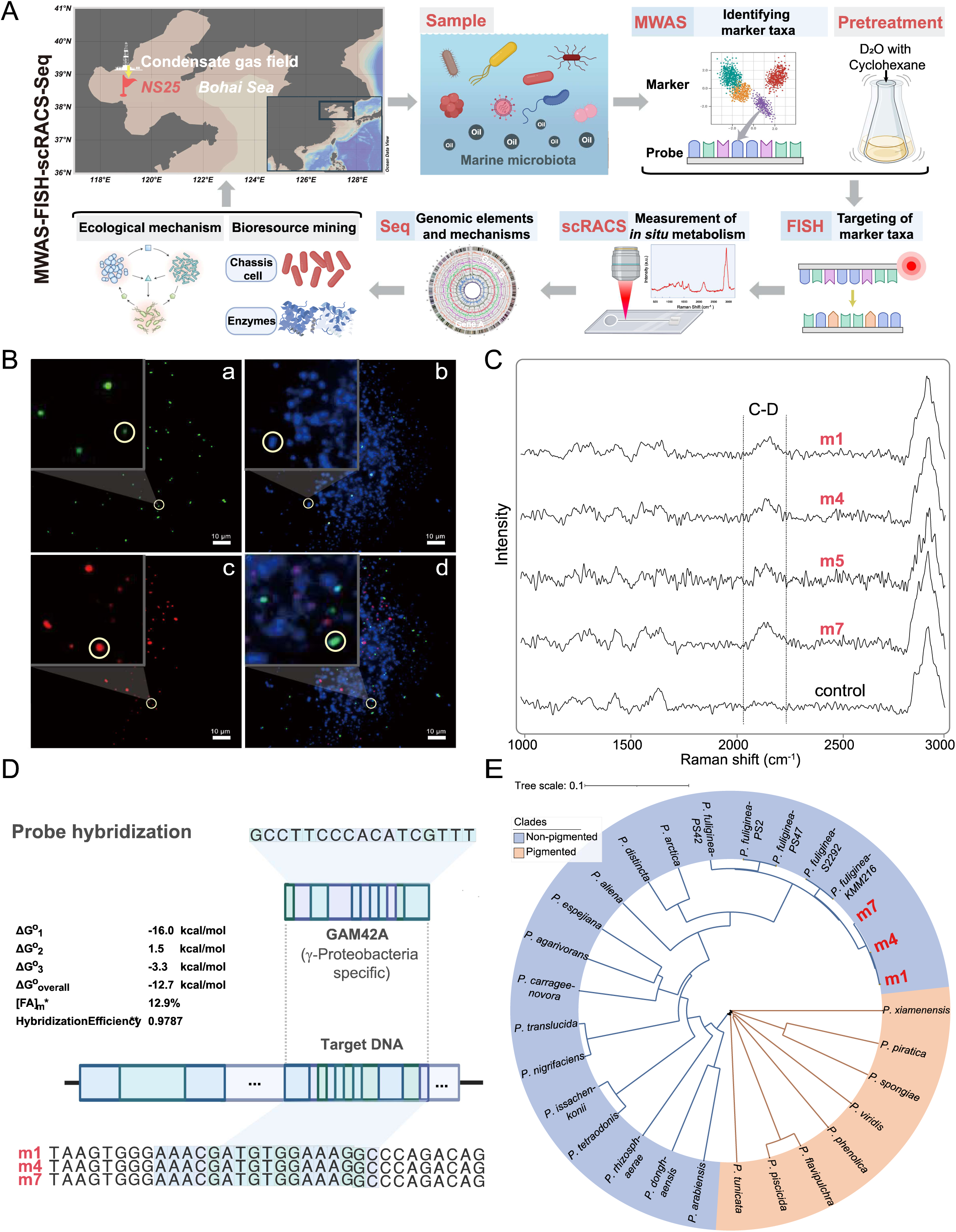
Application of FISH-scRACS-Seq on identifying, sorting and sequencing cycloalkane-degrading γ-Proteobacteria in cycloalkane-contaminated seawater samples. (**A**) The overall strategy of “MWAS-FISH-scRACS-Seq”. (**B**) Photomicrographs of γ-Proteobacteria targeted by CARD-FISH probe from marine microbiota. Identical microscopic fields are displayed for each series. Panel (a) shows γ-Proteobacteria were detected using GAM42a probe; Panel (b) and Panel (c) depict corresponding DAPI staining and autofluorescence of marine bacteria, respectively; Panel (d) displays overlay images of probe signal (green), DAPI staining (blue), and autofluorescence (red); green corresponds to γ-Proteobacteria labelled with Alexa488 using CARD-FISH; blue represents DAPI staining, and red indicates autofluorescence of marine bacteria. Cells appear magenta because of an overlay of DAPI staining and autofluorescence. Scale bar, 10 μm. (**C**) SCRS of the target cells in seawater sample, which were sorted via both “taxon-specific” and “metabolism-specific” features for single-cell genomes. (**D**) FISH-probe-containing genomic regions recovered from SAGs. (**E**) Phylogenetic tree constructed using UPGMA method based on ANI matrix of SAGs and *Pseudoalteromonas* spp. genomes collected from NCBI RefSeq database.

To solve this puzzle, we developed a MWAS-coupled FISH-scRACS-Seq workflow, by employing γ-Proteobacteria-targeted CARD-FISH probes to rapidly identify individual cells shown by MWAS to be associated with cycloalkane degrading, and then specifically profiling their cycloalkane degrading activity *in situ* based on D_2_O-intake rate of a cell (which indicates such activity when cycloalkane is the only carbon source available) via the C-D band in SCRS. The phylogeny-metabolism dual-targeted cells were then sorted and sequenced at one-cell resolution, to mine the pathways and genes responsible for the function (**Fig. 5A**).

Specifically, the seawater samples were incubated with cyclohexane (as sole source of carbon and energy source) plus 50% D_2_O (for tracking metabolic vitality of microbes) at 10 ℃ (temperature of the sampled ocean site; **Methods**). A change in phylogenetic profile of microbiota before and after the cyclohexane treatment is prominent as suggested by 16S rRNA amplicon sequencing, supporting the association of γ*-*Proteobacteria with cyclohexane utilization (**Fig. S4**; **File S1**). When the dissolved oxygen (DO) level was reduced to 0 μM by microbial hydrocarbon respiration, the marine microbiota was sampled to undergo the FISH procedure that employs the GAM42a probe pair which is specific to γ*-*Proteobacteria (**Fig. 5A**; **Methods**). A proportion of cells were successfully labeled, suggesting the presence of a considerable population of γ*-*Proteobacteria, which serve as the basis for subsequent phylogenetically directed screening of metabolic function via SCRS (**Fig. 5B**).

To identify those γ*-*Proteobacteria cells that are degrading cycloalkane *in situ*, the cells with fluorescent signals underwent SCRS acquisition, and those showing C-D bands in SCRS (indicating cycloalkane degrading activity) were sorted, lysed, genome amplified and sequenced via FISH-scRACS-Seq in a one-cell-one-tube manner (**Fig. 5C**; with one cell-free sample as the negative control in each batch). Three one-cell MDA products each with clear MDA bands and positive 16S rRNA PCR results were chosen for 16S rDNA and whole-genome sequencing, producing ∼3 Gb of raw sequencing data for each cell (m1, m4 and m7; **Table S3**). For each cell, GC% of the assembled contigs (> 200 bp; after decontamination; **Methods**) exhibits a normal distribution (**Fig. S3B**), consistent with a clean assembly. Based on lineage-specific marker genes (*42*), 94.35%, 40.49% and 70.88% genome fractions were recovered for m1, m4 and m7, respectively (**Table 2**). Notably, the genomic regions targeted by the CARD-FISH probes were also recovered from each SAG, showing hybridization efficiency of up to 98% (estimated by mathFISH (*59*)) (**Fig. 5D**).

Based on Genome Taxonomy Database (GTDB), the SAGs of m1, m4 and m7 were all classified as *Pseudoalteromonas fuliginea*. Sequence comparison with publicly available *Pseudoalteromonas* genomes via average nucleotide identity (ANI; **Fig. 5E**; **Methods**) revealed >97% similarities of the one-cell *P. fuliginea* genomes with reference genomes, and >99.5% similarities among m1, m4 and m7. To test the value of FISH-scRACS-Seq in reconstructing genomes of the individual cyclohexane-degrading cells, we compared these three SAGs to the seven *P. fuliginea* MAGs aforementioned. (*i*) Most of the MAG-specific genes (versus SAGs) are from two contigs that were either mis-assembled or mis-binned (**Fig. 6A**), pointing to errors in MAG. (*ii*) In SAG m1, four SAG-unique genes are detected which are supported by metagenomic sequencing reads but not recovered in the MAGs (**Fig. 6B**), suggesting the importance of higher gene coverage in SAG. (*iii*) Comparison of core-genome SNPs between SAG m1 and MAGs (based on high-quality SNPs; **Methods**) reveals many cell-specific mutations in the SAG that are absent from the MAGs, with some carrying biological consequence such as introduction of premature stop codons that lead to unstable or nonfunctional proteins (**Fig. 6C**; **Fig. S5**). (*iv*) The SAGs recover many more insertion sequences (ISs) than MAGs (in percent of genomic length, 0.78% vs 0.36%; Wilcoxon test, *p* < 0.05; **Fig. S6**). Therefore, the FISH-scRACS-Seq derived SAGs more completely and accurately reconstruct the genomes of individual cyclohexane-degrading cells.

**Figure 6.**
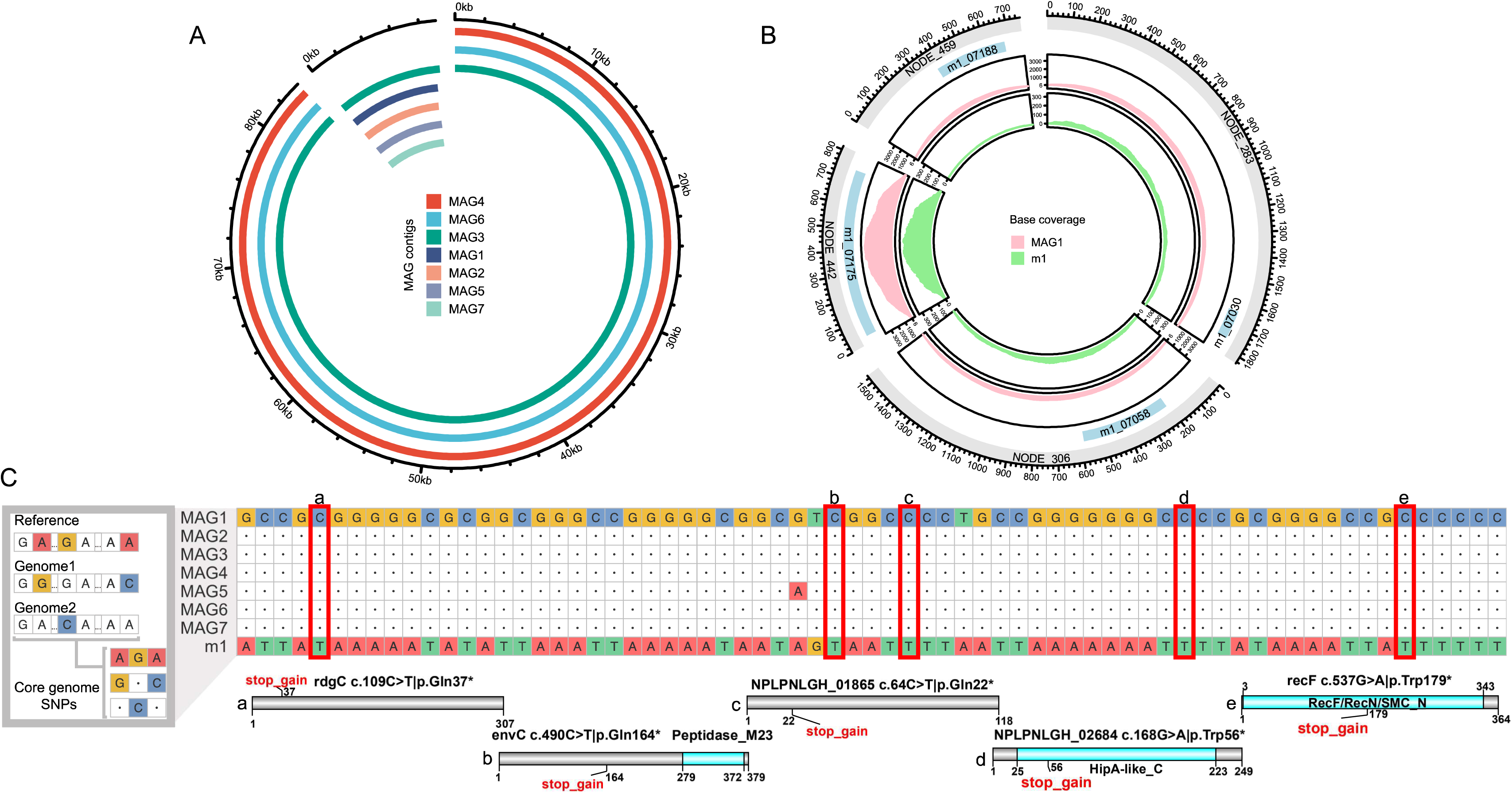
Comparison between the FISH-scRACS-Seq derived SAGs and the shotgun sequencing derived MAGs from cyclohexane-contaminated seawater. (**A**) MAGs recovered two contigs from unknown source, which are not included in FISH-scRACS-Seq derived SAGs. (**B**) FISH-scRACS-Seq derived SAGs reconstructed four unique genes which have support in reads from MAGs but cannot be successfully assembled and binned. (**C**) Core-genome SNP profiling analysis reveals that FISH-scRACS-Seq derived SAGs retain within population genomic heterogeneity. High quality SNPs in m1 SAG were compared to MAGs, with “high impact” SNPs (“stop_gain”: SNP introduces a premature stop codon; shortened polypeptides were also shown) highlighted in rectangular boxes.

Based on these SAGs of *P. fuliginea* that degrade cyclohexane *in situ*, annotation of carbohydrate-active enzymes (CAZymes; via dbCAN2a (*60*)) produced a global view of the glycobiome that potentially underpins the cellular function. In m1, 91 CAZyme genes were identified which suggest the ability to utilize various carbohydrates including pectin, galactoside, glucan, peptidoglycan, chitin, trehalose, porphyran, agarose, alginate, etc (**File S2**). To probe the selection force that shapes these cyclohexane-degrading cells, dN/dS of 2,748 single-copy ortholog genes between m1 and m7 were calculated (*61*). The genes under the strongest positive selection, i.e., with “dN != 0 and dS = 0” and considered “cataclysmic” (225, accounting for 8.2%), are enriched for CAZymes (11 out of 91, **Table S4**; 12% vs. ∼5% in permutations; *p* < 0.05). Strikingly, an alginate lyase harbors four amino acid mutations between m1 (m1_02445) and m7 (m7_06510), suggesting a strong ecological pressure for these organisms to adapt to different types of carbon sources, including but perhaps not limited to alkanes.

### A cytochrome P450_PsFu_ from an *in situ* cycloalkane-degrading *Pseudoalteromonas fuliginea* cell catalyzes cyclohexane degradation

At present, it was not known that members of *Pseudoalteromonas* spp. can degrade cyclohexane, however each of the three *P. fuliginea* cells exhibits strong metabolic vitality *in situ* with cycloalkane as the only carbon source (**Fig. 5C**). To investigate how this occurs, we focused on the 4.42 Mb, 3,853-gene m1 genome which is of the highest completeness (94.35%; **Table 2**; **Table S3**). In m1, a three-component cytochrome P450 systems (class I/class B) was discovered (also in m7; **Fig. 7A**), which consists of *yji*B (“m1_05218”; encoding a cytochrome P450 protein of P450_PsFu_; **Fig. 7A**), *cam*A (“m1_05216”; putidaredoxin reductase) and *fdx*E (“m1_05217”; ferredoxin).

**Figure 7.**
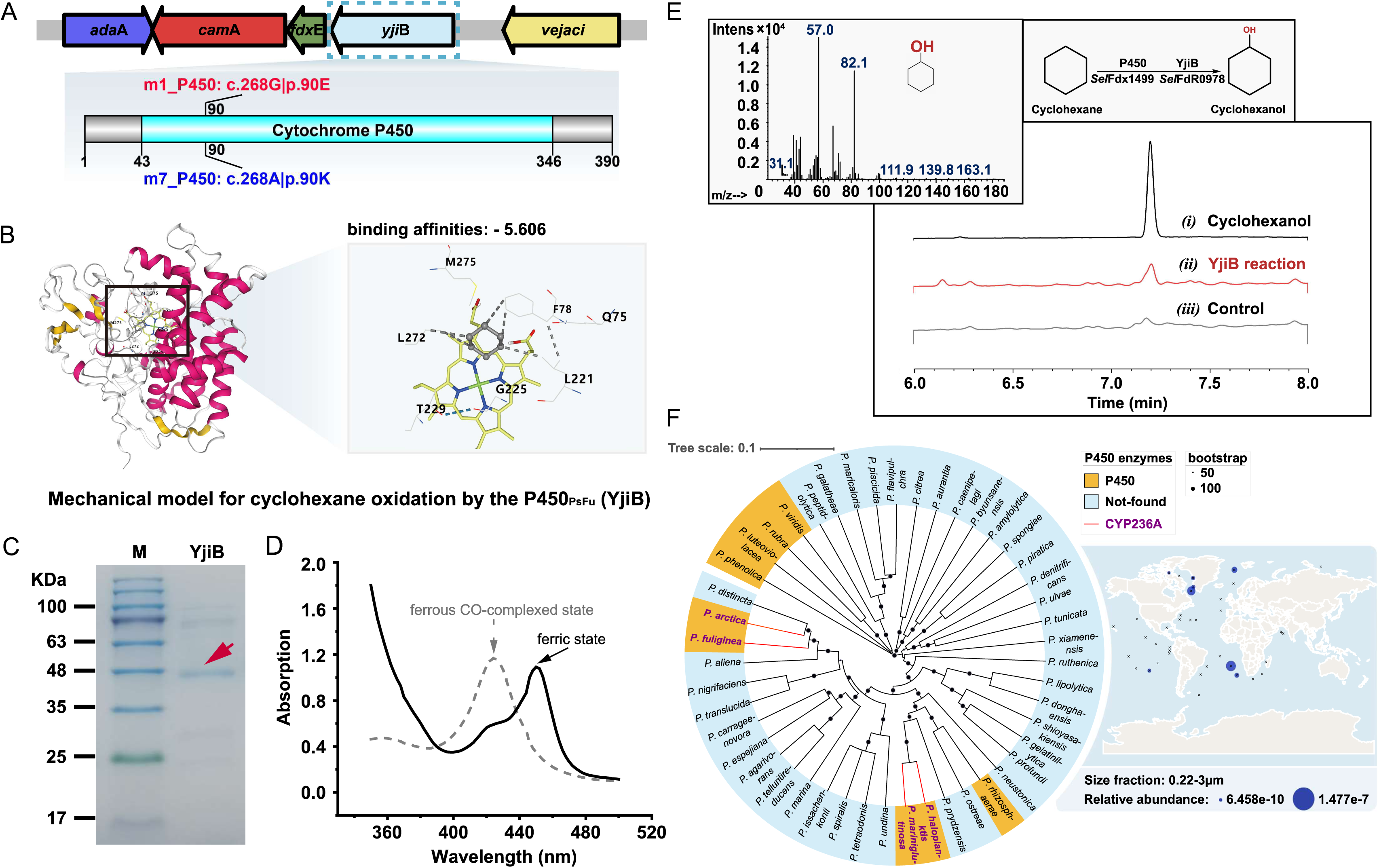
FISH-scRACS-Seq based discovery and experimental validation of cytochrome P450_PsFu_ from *Pseudoalteromonas fuliginea* that catalyses cyclohexane degradation in cyclohexane-contaminated seawater. (**A**) A three-component cytochrome P450 system was recovered in FISH-scRACS-Seq derived SAGs of m1 and m7. The *yji*B gene encodes a cytochrome P450 protein. Genes of *cam*A, *fdx*E and *yji*B constitute three component P450 system. There is one amino acid mutation at position 90 for m7_P450. (**B**) Molecular docking analysis support P450_PsFu_ as cyclohexane degrading enzyme. The protein-ligand docking simulation reveals that cyclohexane (colored in gray) can conjunct with P450_PsFu_. The potential active sites of cyclohexane to P450_PsFu_ are marked. (**C**) SDS-PAGE analysis of purified cytochrome P450_PsFu_. (**D**) CO-bound reduced difference spectra of cytochrome P450_PsFu_. Dotted line, absorbance spectra of cytochrome P450_PsFu_ in ferrous CO-complexed state. Solid line: absorbance spectra of P450 in ferric state. This assay was also used to determine the concentration of functional cytochrome P450 enzyme using the extinction coefficient of 91,000 M^-1^ cm^-1^. (**E**) GC chromatograms and GC-MS of product from the P450_PsFu_-mediated reaction. (*i*) Authentic standard of cyclohexanol; (*ii*) Reaction of cyclohexane with YjiB in the presence of *Sel*FdR0978 and *Sel*Fdx1499; (*iii*) Negative control of (*ii*) with boiled YjiB. (**F**) CYP236A enzymes found in reference genomes of the *Pseudoalteromonas* genus (left panel) and global marine ecosystems (right panel). Among the reference genomes of 47 *Pseudoalteromonas* species, only four harbor CYP236A P450 subfamily genes. At global oceans, CYP236A enzymes are of low abundance and mostly found in colder seas.

In protein sequence, P450_PsFu_ is of very low similarity (25.8%) to the CYP450cha from *Acidovorax* spp. (AKJ87746.1 (*62*)) which is the only P450 known to transform cyclohexane to cyclohexanol so far. In fact, P450_PsFu_ belongs to the CYP236A subfamily which reportedly uses 6-*O*-methyl-D-galactose (G6Me; an abundant monosaccharide of algal agarose and porphyrin) as substrate (*63*). However, considering the promiscuity of substrate specificity in P450 enzymes (*64, 65*), we hypothesize that P450_PsFu_ can oxidize cyclohexane. To test this hypothesis, we started by molecular docking to probe the affinity between P450_PsFu_ and cyclohexane (**Methods**; **Fig. 7B**), which reveals that cyclohexane can bind to P450_PsFu_ through hydrogen bonds and strong electrostatic interactions, and the affinity is actually even higher than that to CYP450cha (i.e., the positive control; **Fig. S7**). Intriguingly, the binding sites/residues of cyclohexane in P450_PsFu_ are identical to the CYP236A subfamily protein of P450_ZoGa_ (**Fig. S8**). In fact, most of the binding sites/residues of cyclohexane/G6Me are conserved across CYP236A-family proteins (**Fig. S9**). Thus, the CYP236A subfamily including P450_PsFu_ may be a previously unknown group of P450 monooxygenases for cyclohexane.

To validate this hypothesis, we conducted an *in vitr*o enzyme activity assay. The sequences of P450_PsFu_, plus its native redox partners of *cam*A, and *fdx*E, were optimized based on *Escherichia coli* codon preferences, and then expressed and purified to homogeneity (**Fig. 7C**). The CO-bound reduced difference spectra of P450_PsFu_ protein display a characteristic peak at 450 nm, confirming the expression of functional P450 enzymes (**Fig. 7D**). As expression of CamA and FdxE in *E. coli* was unsuccessful, *Sel*FdR0978 and *Sel*Fdx1499 from the cyanobacterium *S. elongatus* PCC 7942 was employed as surrogate redox partner proteins instead, to reconstitute the *in vitro* activity of P450_PsFu_ (**Fig. 7E**). The absorption spectra of *Sel*FdR0978 displayed characteristic peaks at around 456 nm and a shoulder at 396 nm, indicative of functional FAD enzymes. *Sel*Fdx1499 exhibited the typical UV-Vis spectra with maximum peaks at approximately 420 nm, which are characteristic of UV-Vis spectra for proteins containing Fe_2_S_2_ cluster (**Fig. S10**). Thus, these two surrogate redox partners are functionally active.

An *in vitro* enzymatic assay for P450_PsFu_ activity was therefore established (**Methods**). Notably, as the low efficiency of NAD(P)H coupling is frequently a significant constraint on the activity of a reconstituted P450 system (due to the extra drain of NAD(P)H during reaction (*66*)), we employed an NADPH regeneration system based on GDH/glucose, in order to detect the P450 activity. GC-MS analysis of the product profiles of reaction reveal a compound with retention time and ionized fragments identical to those of the pure cyclohexanol (**Fig. 7E**). Thus, P450_PsFu_ is able to convert cyclohexane to cyclohexanol, which is the first and the rate-limiting step of cyclohexane degradation, with the support of *Sel*FdR0978 and *Sel*Fdx1499.

Although members of the CYP236A P450 subfamily such as P450_FoAg_ and P450_ZoGa_ can oxidize the methyl group on G6Me (*63*), it is much more difficult to oxidize the inert hydrocarbon bonds, due to their much higher reaction energy requirements. Thus, the metabolic activities of P450_PsFu_ enzyme and of a *Pseudoalteromonas* spp. in oxidizing cyclohexane on the cycloalkyl group are new and surprising. These findings also reveal the diverse substrate specificity, and the ecological versatility (i.e., in macromolecule degradation), of the CYP236A P450 subfamily.

### Ecological significance of the newly discovered cycloalkane monooxygenase of P450_PsFu_

The CYP236A P450 subfamily enzymes are predominantly found in Bacteroidetes and γ-Proteobacteria (*63*). Among the 47 sequenced *Pseudoalteromonas* spp. genomes, only four species that are distributed across multiple organismal branches harbor members of this subfamily (**Fig. 7F**). Thus, although all located on chromosomes and not in plasmids, these genes are “accessory” genes found in only a very small set of genomes such as a few *Pseudoalteromonas* spp. and likely originated via horizontal gene transfer or independent evolution (*63*). Two of the four species (*P. fuliginea* PS2 and *P. mariniglutinosa* NCIMB 1770) were originally isolated from surface of marine algae, suggesting a symbiotic relationship underpinned by P450-mediated bacterial utilization of algal polysaccharides. Interestingly, although P450_PsFu_ is discovered in a *P. fuliginea* from the Bohai Sea at the northeastern Pacific Ocean, another of the species of *P. arctica* A 37-1-2 was isolated from 4 ℃ seawater of the Arctic Sea in Spitzbergen (Norway), a location rich in oil resources and having encountered oil spills (*67, 68*). Therefore, cyclohexane degradation by the CYP236A subfamily of P450 enzymes is likely of ecological significance at a global scale.

From the KMAP metagenomic database which includes MAG-derived genes of diverse environments (including marine, soil, rhizosphere, lake, wastewater, saltmarsh, etc (*69, 70*)), a search for CYP236A P450 subfamily enzymes revealed only eight such genes, and they all originated from seawater, marine sediment or freshwater (**Table S5**). Moreover, from the Ocean Microbial Reference Catalog v2 (OM-RGC.v2; (*71*)) and Ocean Gene Atlas v2.0 ^116^ (**Fig. 7F**; Methods), 32 genes encoding such enzymes were found in 22 Tara Oceans projects (**Table S6**), with 20 of the 22 projects being from the Arctic Ocean and with temperatures of sampled sites ranging from -1.5 to 8.5 ℃. The predominant distribution of these enzymes in low-temperature marine ecosystems is likely consistent with that of oceanic cyclohexane sink (*72, 73*).

Collectively, these results suggest the CYP236A subfamily of P450 enzymes as represented by P450_PsFu_ are enzymes that are numerically rare in their microcosms and spatially constrained to specific environments, but likely have contributed to cyclohexane degradation at low-temperature oceans on a global scale. These ecological features can be exploited for bioaugmented removal of hydrocarbon pollutants during oil spills.

## Discussion

In many ecosystems, crucial functions can be mediated by numerically rare members of the microbiota that are not yet cultured (*74, 75*), yet validation of such roles and mining the underlying pathways or enzymes are usually difficult, due to the inability to profile their functions and the corresponding genomes *in situ*. One such example is cyclohexane biodegradation in contaminated seawater from a condensate gas field at the Bohai Sea. Although MWAS that correlates microbiota structure with cyclohexane level has identified the group of γ-Proteobacteria as marker organisms for this crucial ecosystem trait (*55*), these functional cells remain uncultured and their functional genes are too low in relative abundance to be detected by WGS of the microbiota. By establishing FISH-scRACS-Seq and coupling it to MWAS, we were able to, in a phylogeny and metabolism dual-directed manner, trace the *in situ* cyclohexane activity to individual *P. fuliginea* cell and then further to P450_PsFu_ which represents a previously unknown group of cyclohexane monooxygenases, and of cyclohexane-degrading genus.

The phylogeny directedness of MWAS-derived FISH probes ensures the metabolic activities of cells (and pathways and enzymes encoded by their genomes) discovered by FISH-scRACS-Seq are of ecological relevance. On the other hand, the metabolism directedness of scRACS-Seq ensures the individual cells recovered are actually performing the right function *in situ* (which is critical as it is well-known that genome-based metabolic reconstruction or functional assays in pure culture are unable to reliably predict the *in situ* function of a cell in a microbiota at its native state (*18, 76*)). In a natural microbiota sample, cells with an interested metabolic activity that is nonrelevant in ecosystem functioning are abundant, so are those from a target taxonomy yet not performing the target biodegradation activity. Therefore, the dual directedness of FISH-scRACS-Seq greatly elevates the efficiency in dissecting “Who is doing What” in a microbiota, particularly for the low-abundance yet functional members of microbiota, such as the *P. fuliginea* cell and the P450_PsFu_.

Cyclic alkanes, common in hydrocarbon reservoirs and gas condensates, are highly toxic to aquatic life and pose an ecological risk during oil spills, while the mechanisms of biodegradation in marine ecosystems remain poorly understood (*55*). In particular, cyclohexane, a common organic compound found in various industrial and environmental settings, is biotoxic yet its biodegradation is difficult due to its chemical structure (*77*). In cyclohexane degradation, its oxidation to the non-biotoxic cyclohexanol represents the first and the rate-limiting step. The discoveries of *P. fuliginea in situ* and of P450_PsFu_ for this crucial step are unexpected, as no *Pseudoalteromonas* spp. or members of the CYP236A P450 subfamily were known to have this talent. Notably, as the *P. fuliginea* genome lacks enzymes for subsequent utilization of cyclohexanol, which however is of much higher bioavailability to microbial degraders than cyclohexane, it is possible that *P. fuliginea* is a keystone species at the start of a food chain and collaborates with other symbiotic bacteria for the complete mineralization of cyclohexane. Therefore, although numerically rare in their microcosms and spatially constrained to specific environments, they might have contributed to cyclohexane degradation in low-temperature oceans at a global scale.

Further development of FISH-scRACS-Seq can take multiple directions. (*i*) Although the use of CARD-FISH probes improves the detection of target cells (*78*), multiplexing of probe hybridization is possible, which should allow simultaneous interrogation of phenome-genome-gene links for multiple marker organisms (*79, 80*). Moreover, in addition to taxonomical markers, functional genes can also be targeted by a FISH probe via nucleotide sequence, therefore, FISH-scRACS-Seq can be extended to dissect a target gene’s *in vivo* function in mutant libraries, microbiota, or even plant or animal tissues. (*ii*) In microbiota RACS-Seq, by creating a RAGE chip that preserves cell vitality (*31*) and designing a Hotja Phi29 enzyme that reduces bias in DNA amplification in one-cell MDA reactions (*81*), we have demonstrated production of high-quality SCRS plus corresponding high-coverage genomes at precisely one-cell resolution directly from diverse ecosystems such as urine (*31*), gastric biopsy (*29*), soil (*24*), seawater (*25*), wastewater (*32*) and probiotics products (*33*). However, at 3∼8 cells/min (*31*), the sorting throughput should be elevated, e.g., by improving RAGE-chip design (*82*) and incorporating A.I.-based image analysis and automation into the RAGE operation (*83*). Flow-mode RACS systems that sort at much higher throughput can also be adopted (*84-86*), to enable much deeper sampling of complex microbiota for a diverse set of cellular functions via SCRS (*11, 18*). (*iii*) Cultivation of the target cells after FISH-scRACS operation is highly desirable, particularly since cells can remain viable after RACS, as demonstrated in scRACS-Culture for phosphate-solubilizing bacteria in wastewater (*32*), and pool-based RACS-Culture for mucin-degrading microbes from mouse colon microbiota (*76*). As chemical cross-linking or fixation with paraformaldehyde (to stabilize the cells and partial cell wall lysis) and ethanol (to enable probe penetration) would cause cell death, live-FISH techniques followed by RACS-Seq should be explored that allows bacteria to survive the FISH procedure (*37*).

In summary, mechanistic dissections of microbiota function have greatly lagged behind the explosive pace of MWAS generating marker organisms (and marker genes) for ecosystem traits. FISH-scRACS-Seq can bridge this long-standing gap by efficiently unveiling enzymes, pathways, genomes and *in situ* metabolic functions specifically targeting those cells revealed by MWAS as of ecological relevance, regardless of their cultivability. Therefore, we propose the complete MWAS-FISH-scRACS-Seq (e.g., **Fig. 5A**) as a rational and generally applicable platform to systematically and thoroughly dissect and mine microbiota function from the pletheora of ecosystems on Earth.

## Materials and Methods

### Experimental Design

The FISH-scRACS-Seq workflow consists of three steps (**Fig. 1**). In Step 1 (i.e., “FISH”), individual cells of a target taxon are directly localized in a microbiota sample, via a taxon-specific FISH probe. In Step 2 (i.e., “scRACS”), post-FISH cells are distinguished and sorted based on not just the target phylogeny (via the FISH probe) but also the target metabolic phenome (via the SCRS). In Step 3 (i.e., “Seq”), the post-FISH-RACS cells in droplets would undergo cell lysis, Multiple Displacement Amplification (MDA), and genome sequencing in an indexed, one-cell-one-tube manner In this way, specifically for those cells of target phylogeny in a microbiota, the target metabolic activity *in situ* is directly traced to genome sequence at single-cell resolution.

### Bacterial species, media and growth conditions

The series of mock microbiota include the three bacteria of *Escherichia coli* K-12 DH5α, *Micrococcus luteus* D11 (isolated from soil environment in our laboratory) and *Bacillus subtilis* H6, and the one fungus of *Saccharomyces cerevisia*e BY4742. Each of the strains was grown in pure culture. *E. coli* K-12 DH5α, *M. luteus* D11 and *B. subtilis* H6 were all cultured in Luria-Bertani (LB) medium (Tryptone, Yeast extract, NaCl, pH 7.0) and incubated at 37 ℃. These three strains were diluted to an OD 600 of ∼0.5 and inoculated at a ratio of 1:10 into 4 mL of LB medium, respectively. *S. cerevisiae* BY4742 were cultured in YPD medium (Yeast Extract, Peptone, glucose, pH 6.5∼6.8) and incubated at 30 ℃. *S. cerevisiae* BY4742 was diluted to an OD 600 of ∼0.5 and inoculated at a ratio of 1:10 into 4 mL of YPD medium. In these experiments, we assumed a constant relationship between OD and cell concentration for all the species.

For deuterium isotope labeling, 50% D_2_O (vol/vol) (99.9 atom% D, Sigma-Aldrich, Canada) was used in all the above media. To prepare the media for deuterium isotope labeling, 2×medium was prepared with water and autoclaved, and then diluted to 1×medium with filtered pure D_2_O, so that the eventual level of D_2_O is 50%. Each of the microorganisms was incubated in the respective medium containing 50% D_2_O until reaching the logarithmic phase, washed using distilled water, and mixed to form the synthetic consortia with defined structure. The mock microbiota were then subjected to CARD-FISH labeling, single-cell Raman spectroscopy and SCRS-based sorting, respectively.

### Probing the active cycloalkane-degraders in situ by D_2_O labelling

A bottom water sample of NS25 was collected from a large condensate gas field in the Bohai Sea during the spring of 2020. NS25T18 was derived from NS25 by enriching cycloalkane-degrading activities via incubating the seawater with methylcyclohexane as described (*55*). Cyclohexane and D_2_O were then added to NS25T18 to probe the metabolically active cycloalkane degraders. Glass serum bottles (CNW, 10 mL) outfitted with an oxygen sensor spot (Pyroscience, OXSP5) were used in the incubation to allow for contactless monitoring of oxygen concentration. Specifically, 2 mL of ONR7a, 4 mL of D_2_O, 2 mL of bloomed culture NS25T18, and 2 µL of cyclohexane (Aladdin, purity 99.5%) were added into the sterile glass serum bottles. As controls, 2 mL of ONR7a, 4 mL of H_2_O, 2 mL of bloomed culture, and 2 µL of cyclohexane were added into the sterile glass serum bottles. Two duplicates were prepared for each treatment.

### Extracting bacteria from soil and D_2_O labeling of microbial cells

To extract the cells from soil, the soil slurries generated by adding 1 g soil into 5 mL 1× PBS buffer supplemented with 25 µL Tween 20 were vortexed for 30 min to free the particle-associated cells. In a new 15 mL centrifuge tube, 5 mL Nycodenz iohexol (1.42 g/mL; Aladdin, China) was added, then the aforementioned supernatant from soil slurries was slowly added to the top of Nycodenz. The tubes were centrifuged at 14,000×g for 90 min at 4 °C with slow acceleration and deceleration. At the middle layer which is between the clear PBS layer and the debris layer, a faint whitish band containing bacterial cells would emerge (*17, 87*). This band was recovered and transferred into a new 1.5 mL Eppendorf tube with a pipette. Then 1 mL ddH_2_O was added to resuspend the cells, and the cells were pelleted by centrifugation at 10,000 ×g for 10 min at 4 °C for 3 times. Finally, the cell pellets were resuspended in 0.2 mL ddH_2_O, which represent the “soil cell extracts”.

As for D_2_O-probed SCRS acquisition and scRAGE-Seq experiments for metabolically active cells, the soil cell extracts were then incubated in PBS with final D_2_O level of 50% at room temperature for 24h based on the previous experimental results (*24*). After that, this D_2_O-labeled cell extracts were subjected to the CARD-FISH procedure.

### The CARD-FISH procedure of microbiota samples

#### Sample preparation and fixation

The aforementioned D_2_O-labelled-cells from pure cultures or environmental samples (i.e., soil and seawater) were centrifuged and fixed in 4% paraformaldehyde (PFA) solution in 1 x phosphate buffered saline (PBS) buffer, respectively. Fixed cells can be stored in 1 x PBS/ethanol (EtOH) at -20 ℃ for up to one month without apparent effect on the hybridization. To inactivate endogenous peroxidases, the fixtures were dehydrated in different concentrations of ethanol (50%, 80%, 100% [v/v] in H_2_O) successively, and then treated in 0.15% H_2_O_2_ in methanol solution for 30 min at room temperature. For permeabilization of cell walls, cells were rinsed with PBS and then consecutively incubated in lysozyme solution (10 mg·mL^-1^ in 0.5 M EDTA, 1 M Tris·HCl (pH 8.0)) for 60 min at 37 ℃. To minimize cell loss during cell wall permeabilization, low-speed centrifugation (6000 rpm, 5 min) was implemented instead of filtration and agarose embedding for cell collection, and specifically, care was taken to avoid excessive agitation during the washing steps. After two washing steps with sterile H_2_O_MQ_, the samples were rinsed with ethanol dehydrated, dried at room temperature, and subsequently stored at -20 ℃ until further processing.

#### Hybridizations with oligonucleotide probes

For CARD-FISH, the 16S rRNA targeted oligonucleotide probes GAM42a (**Table S1**, TAKARA Bio Inc., Japan) was applied to the identification of γ-Proteobacteria. Probe was labeled by horseradish peroxidase (HRP) and Alexa Fluor 488 (Invitrogen, USA) was used to synthesize the tyramide conjugate. Negative controls were performed by probe NONEUB (**Table S1**, TAKARA Bio Inc., Japan).

Hybridizations were performed at 46 °C for 2 h using 300 µL of hybridization buffer (0.9 M NaCl, 20 mM Tris·HCl (pH 8.0), 20% [w/v] dextran sulfate (Sigma Aldrich), 0.02% [w/v] sodium dodecyl sulphate (SDS), 1% [w/v] blocking reagent (Roche, Germany), and 20% [v/v] formamide) together with 1μL probe (50 ng μL^-1^). A negative control, i.e., that without adding the probe, was included. After adding the pre-warmed washing buffer (0.215 M NaCl, 5 mM EDTA (pH 8.0), 20 mM Tris·HCl (pH 8.0), and 0.01% [w/v] SDS), the samples were incubated at 48 °C for 10 min and then equilibrated in 1 x PBS solution for 15 min at RT.

#### Tyramide signal amplification and microscopic evaluation

For tyramide signal amplification in the CARD-FISH experiment, the samples were collected at 6,000 rpm for 5min to remove excess buffer and resuspended in 1mL amplification buffer (2 M NaCl, 0.1% [w/v] blocking reagent, 10% [w/v] dextran sulfate, 0.0015% H_2_O_2_ in 1 x PBS (pH 7.4)), mixed with 1 µL of AF_488_-labeled tyramide solution (1 mg·mL^-1^), and then incubated at 37 ℃ for 30 min in dark. After incubation, the samples were transferred to 1x PBS and soaked in dark for 15 min, followed by a thorough wash with H_2_O_MQ_. Then CARD-FISH preparations were counterstained with 4,6-diamidino-2-phenylindole (DAPI) (1 μg·mL^-1^), subsequently washed in H_2_O_MQ_, dehydrated in ethanol, and air-dried in the dark. The obtained samples mounted with Citifluor (Citifluor Ltd, UK) can be stored at -20 ℃ for up to one month without loss of fluorescence intensity. Microscopic preview was performed by an epifluorescence microscope with a U-RFL-T light source (BX53F2; Olympus, Japan).

In addition, for mock microbiota, cell counts for the target cells were performed by hemocytometer before and after CARD-FISH labeling, respectively, for comparing the change in the proportion of target bacteria before and after CARD-FISH labeling. Briefly, the target bacterial suspension was diluted in an appropriate gradient and subsequently allowed to flow automatically into the counting chamber along the edge of the coverslip, and the quadrant and center compartments of the counting chamber were selected for cell count estimation (*88*).

Experimental details for target-cell sorting, multiple displacement amplification, library construction and next generation sequencing are provided in **Supplementary Materials**.

### Sequencing data analysis

#### Metagenomic data analysis

For the analysis of 16S rRNA gene data, the software package of QIIME2 (version 2022.8) was employed (*89*). The forward and reverse read pairs were denoised, dereplicated and chimera filtered using DADA2 algorithm (*90*) (with default parameters), then clustered based on Vsearch algorithm (*91, 92*) with Greengenes database v13.8 as reference (percent identify cutoff of 97%). At the phylum, class, order, family, genus, and species levels, the relative abundances of the bacterial taxa were calculated and compared, respectively. Finally, biomarkers were discovered using LEFSe at the class and the genus levels (*93*). For analysis of WGS data, sequencing reads were quality controlled, assembled and binned using MetaWRAP pipeline (*94*). Then, all the bins (MAGs) were annotated using Prokka (*95*) and eggNOG-mapper (*96*). The depth of coverage for each contig was obtained by remapping sequencing reads to each MAG using Bowtie2 (*97*) and winding up using BEDTools (*98*).

#### One-cell genome sequencing data analysis

An integrated computational pipeline (SCGS; https://github.com/gongyh/nf-core-scgs) for analyzing single-cell amplified genomes (SAGs) datasets was applied. Briefly, raw sequencing reads were quality controlled using Trim Galore (*99*) in paired end mode for each sample. Then, clean reads were assembled into contigs using SPAdes (*100*) in single-cell mode. Taxonomic composition of assembled contigs (longer than 200 bp) was visualized using BlobTools (*101*). Assembled genomes were annotated using Prokka (*95*), KofamKOALA (*102*) and eggNOG-mapper (*96*). Considering the possibility of DNA contamination for environmental samples, assembled contigs were further split into bins by taxonomic annotations (in the genus level) for each SAG, followed by estimation of genome completeness using CheckM (*42*).

#### Heterogeneity analysis

The taxonomy of each SAG was obtained via GTDB-Tk (*103*). The average nucleotide identity (ANI) between each SAG was calculated using OrthoANIu v1.2 (*104*). Orthologs between SAGs/MAGs were identified using OrthoLoger v2.6.2 (*105*). SNP variants between core-genomes of SAGs/MAGs were called using Parsnp v1.7.2 (*106*). High quality SNPs in SAGs were called using MonoVar (*107*) (shared ones were used for subsequent analysis), and annotated using SnpEff v5 (*108*).

#### Comparison of single-cell sequencing performance via SAG

The FISH-scRACS-Seq-derived SAGs from soil were compared to scRAGE-Seq-derived SAGs (*24*) and SAG-gel-derived SAGs (*51*). To be fair, only SAGs from soil and belonging to γ-proteobacterium were considered, and all SAG contigs shorter than 1000 bp were filtered and removed prior to comparison. For scRAGE-Seq, five SAGs (SR9, BSR2, BSR3, BSR5, BSR11) were obtained for comparison. The metrics of FISH-scRACS-Seq-derived SAGs and scRAGE-Seq-derived SAGs were calculated as described before (*24, 31*). For SAG-gel-derived SAGs, published metrics were used.

#### Sequence analysis of CYP236A subfamily proteins

*(i)* A total of 173 proteins that are homologous to P450_PsFu_ were retrieved from Uniref90 database (*109*) using MMseqs2 ((*110*); with parameters of “--min-seq-id 0.4 -c 0.8”). Multiple sequence alignments of those proteins were performed using MAFFT v7.508 (*111*). Residue conservations were visualized using WebLogo v3 (*112*). (*ii*) The sequences of P450_PsFu_ (in SAG m1 and m7), P450_ZoGa_ (WP_013995999.1) and P450_FoAg_ (WP_038530297.1) were compared using MAFFT v7.508 (*111*) and visualized using the R package ggmsa (https://cloud.r-project.org/package=ggmsa). (*iii*) The CYP236A P450 subfamily proteins were searched in the environmental metagenomic databases of KMAP (*69*) and OM-RGC.v2 (*71*) using P450_PsFu_ as the query. Initially, candidate CYP236A proteins were selected based on homology comparisons using blastp (*113*) or diamond blastp (*114*); then, the sequence identities between P450_PsFu_ and all candidate CYP236A proteins were calculated using ggsearch36 (*115*); finally, proteins in the CYP236A P450 subfamily were filtered out (with a criterion of protein identities > 55%) using custom scripts. Phylogenetic trees were visualized using the iTOL online service (*116*). The abundance of CYP236A enzymes on the ocean map was visualized using the web service of Ocean Gene Atlas v2.0 (*117*), with m1_P450 as query sequence and e-value of 1E-160.

#### Molecular docking analysis

Firstly, the three-dimensional structure of candidate P450 enzymes was predicted using ESMFold (*118*). Then, to analyze the binding affinities and modes of interaction between the candidate P450 enzymes and cyclohexane (and ligand Heme), CB-Dock2 was employed for docking calculations and visualizations (*119*), via an online service (https://cadd.labshare.cn/cb-dock2/php/blinddock.php).

### Purification and in vitro enzymatic assay of the cytochrome P450_PsFu_

#### Protein purification

The *yji*B gene which encodes cytochrome P450_PsFu_ was codon-optimized and inserted between the *Nde*I and *Not*I restriction sites of pMAL-c5E to create pMAL-c5E-*yjiB*. This plasmid was transformed into *E. coli* BL21 (DE3). A single colony of the transformant was inoculated into LB medium containing 50 mg/L ampicillin. The seed culture grown overnight was used for 1:100 inoculation of 1 liter of LB medium containing 50 mg/L ampicillin, 1 mM thiamine, 10% glycerol, and rare salt solution at 37°C (*120*). Protein expression was induced with 0.2 mM isopropyl-*β*-d-thiogalactopyranoside (IPTG), and 1 mM *δ*-aminolevulinic acid was added as the heme synthetic precursor until the optical density at 600 nm (OD600) reached 0.6 to 1.0. The cells were then incubated with shaking for 20 h at 18 °C. The protein was purified according to a previously developed procedure (*120*) and digested with enterokinase to release MBP. The native redox partner *cam*A and *fdx*E were unable to express in *E. coli*. Therefore, the frequently-used surrogate redox partner *Sel*Fdx1499/*Sel*FdR0978 from *S. elongatus* PCC 7942 was chosen for analyzing the *in vitro* enzyme activity of YjiB. SDS-PAGE (**Fig. 7C** and **Fig. S10**) verified the purification of all enzymes, which were stored at -80 °C. The UV-visible spectra were obtained on a UV-visible spectrophotometer (Varian, UK). The functional concentration of P450 YjiB was calculated from the CO-bound reduced difference spectrum using an extinction coefficient (ε_450–490_) of 91,000 M^-1^ cm^-1^ (*66*). The concentrations of *sel*Fdx1499 and *sel*FdR0978 were determined by measuring the absorbance at selected wavelengths. The extinction coefficients of *Sel*FdR0978 and *Sel*Fdx1499 were determined to be ε_454_ = 19120 M^-1^ cm^-1^ and ε_460_ = 11270 M^-1^ cm^-1^, respectively (*66*).

#### Enzymatic assay of the cytochrome P450_PsFu_

The standard assay contained 5 μM P450, 50 μM Fdx, 25 μM FdR, 500 μM substrate, 0.5 mM NAD(P)^+^ and NAD(P)H regeneration system (10 mM glucose and 2 U glucose-6-phosphate dehydrogenase) in 200 μL reaction buffer (50 mM potassium phosphate buffer, pH 7.4). A ratio of 1:10:5 for P450/Fdx/FdR was used to ensure adequate electron supplies and efficient P450-mediated conversions. The reactions were incubated at 30 ℃ for 12 h and quenched by adding an equal volume of ethyl acetate and vortexing for organic extraction. After high-speed centrifugation, the organic phases were directly used as samples for GC or GC-MS analysis. GC and GC-MS analyses were performed on an Agilent 7980A gas chromatograph equipped with an Agilent 5975C mass selective detector (Agilent Technologies, Little Falls, DE, USA) and a HP-INNOWAX capillary column (30 m × 0.25mm × 0.25µm, Agilent, USA). The program used for GC-MS analysis was as follows: carrier gas: He, flow rate 1.0 mL/min, no splitting, inlet temperature: 300 °C, the oven temperature was held at 40 °C for 3min, increased to 200 °C at a rate of 15 °C/min, then increased to 240 °C and held for 8 min, transfer line temperature: 280 °C, ion source temperature: 230 °C.

## Acknowledgements

We are grateful to Zongze Shao (from Third Institute of Oceanography, Ministry of Natural Resources of China) for his insightful comments.

## Author Contributions

Conceptualization: XJ, XYJ

Methodology: XYJ, YM, ZDD, JC

Investigation: YM, YSR, YCL, WHS

Visualization: YHG, XYJ, YM

Supervision: YTL, JZ, YTJ, BM, ZQC, SYL

Writing: XJ, XYJ, YHG, LM, YM, ZSC

## Data and materials availability

The sequence data reported in this study have been deposited in the NCBI SRA database under BioProject ID: PRJNA890413, PRJNA891066, PRJNA814381, and PRJNA855277. All data are available in the main text or the supplementary materials.

## Supplemental Figures, Tables and Files

**Figure S1. Agarose gel images of the multiple displacement amplifications (MDAs) and 16S rRNA gene validation processes for target γ-Proteobacteria cells obtained via the FISH-scRACS-Seq.** MDA products and PCR products of the 16S rRNA genes of the target γ-Proteobacteria cells from the pure cultured *Escherichia coli* K-12 DH5α (A) and from a four-species mock microbiota (B-D, three batches of experiments (B, C and D) were performed). Lane NC, empty droplet (i.e., without cells); Lane N, negative control for PCR (i.e., without adding template); Lane P, positive control for PCR. The non-specific amplification of MDA is due to formation of primer dimers in the MDA reaction.

**Figure S2. Validation of the probe GAM42a targeting (A) *Micrococcus luteus* D11 (*Ml, non* γ-Proteobacteria), (B) *Bacillus subtilis* H6 (*Bs, non* γ-Proteobacteria) and (C) one fungus of *Saccharomyces cerevisiae* BY4742 (*Sc, non* γ-Proteobacteria), respectively.** Panel (a) shows the photomicrographs of strain hybridized with HRP-labeled oligonucleotide probes GAM42a; Panel (b) represents DAPI staining (blue) and the same field of view which shows a color combined image recorded by epifluorescence microscopy in Panel (c); Phase-contrast photomicrographs is shown in Panel (d).

**Figure S3. GC distributions of contigs from the FISH-scRACS-Seq derived SAGs.** GC distributions of the contigs from the SAGs that correspond to FISH-scRACS-sorted cells in soil (A) and seawater (B), respectively. Black curves represent GC distribution of recovered draft genomes. A sliding window of 200 bp along each contig was used to extract sequence fragments and then calculate GC contents. Red curves show theoretical normal distribution with similar mean and standard deviation to the corresponding GC distribution. The GC contents of these sets of contigs exhibit normal distribution, supporting the integrity of the one-cell assemblies. (C) Mapping SAG contigs to a plasmid sequence (accession NZ_CP024444.1) reveals the recovery of a plasmid-encoded antimicrobial resistance gene in the SAG of s9.

**Figure S4. The bacterial community structures of the original seawater and the D2O-spiked enrichment cultures derived from the Bohai Sea, respectively.**

**Figure S5. Read mappings to the MAG1 genome showed sensitive and reliable recovery of chromosomal mutations in SAG1**. Positions of the five SNPs which introduce premature stop codons were shown here. Mismatches in read mappings were highlighted using different colors (not gray).

**Figure S6. The insertion sequences (ISs) recovered from bulk sample, SAGs and MAGs, respectively.** Genome sequences of isolated *P. fuliginea* (“Isolate”) are retrieved from NCBI RefSeq database. Single-cell genomes (“Single-cell”) include SAGs of m1, m4 and m7. MAG genomes (“Metagenome”) consist of MAG1-MAG7.

**Figure S7. The protein-ligand docking simulation reveals that cyclohexane (colored in orange) can conjunct with the CYP450cha (AKJ87746.1 from *Acidovorax* spp.).**

**Figure S8. Sequence logo of CYP236 family proteins selected from the UniRef90 database.** The active sites of G6Me to P450ZoGa, which contains all the potential active sites of cyclohexane to P450PsFu, are highlighted in red squares.

**Figure S9. Multiple sequence alignments of P450PsFu (in SAGs m1 and m7), P450ZoGa and P450FoAg.** The active sites of G6Me to P450ZoGa are conserved and marked with black asterisk.

**Figure S10. SDS-PAGE (A) and UV/Vis absorption spectra (B) of *Sel*FdR0978 and *Sel*Fdx1499.**

**Table S1. Probes used in FISH-scRACS-Seq and primers used for PCR amplification of 16S rRNA gene for MDA products.**

**Table S2. Performance of FISH-scRACS-Seq targeting *Escherichia coli* K-12 DH5α (Ec).** The GC content and the genome size of reference genome is 50.79% and ∼4. 64 Mb (RefSeq: NC_000913.3).

**Table S3. Sequencing and assembly statistics for single-cell genomes from the soil and seawater produced by FISH-scRACS-Seq.**

**Table S4. Carbohydrate-active enzymes in sample m1.**

**Table S5. List of CYP236A P450 subfamily enzymes searched from the KMAP metagenomic database.**

**Table S6. List of CYP236A P450 subfamily enzymes searched from the Ocean Microbial**

## Reference Catalog v2

Supplementary file 1. List of the taxonomy abundance for the seawater samples (A, B and C represent three biological replicates).

Supplementary file 2. Genes identified as CAZymes in SAG of m1.

