## Supplementary Materials for "Phylogeny-metabolism dual-directed single-cell genomics for dissecting and mining ecosystem function"

Xiaoyan Jing *et al.*

### Supplementary Text

### Materials and Methods

#### *Detection and isolation of CARD-FISH&$D_2O$-labeled cells by RAGE*

The FISH-scRACS-Seq procedure was performed in a RACS-Seq instrument (Qingdao Single-cell Biotechnology, China). Before Raman test, the CARD-FISH&$D_2O$-labeled bacterial samples were washed to remove residual media, and then resuspended by adding deionized water to dilute for performing the following fluorescent detection, Raman measurement and cell sorting. The prepared bacterial solution (~ 1mL) was hanged up on the sample holder and then loaded into the RAGE chip. The well was filled with mineral oil (2% wt EM90) when the cell phase reached the open well. The sample holder height was adjusted to obtain a balance between the water phase and oil phase. For the fluorescent labeled cells, their Raman spectra were acquired with a modified confocal Raman microscope. 50 × dry objective (NA=0.65, Leica, German) was used for sample signal acquisition and optical tweezers, while a 10× dry objective was used for observation of droplet generation and transportation. The laser was switched to 1,064 nm, to trap and move a single target cell to the edge of the aqueous phase, while clearing away other cells near the tip to ensure single-cell encapsulation. The sample holder was elevated to generate only one droplet, then lowered to the original height, so a single target cell could be isolated and encapsulated within the droplet. The density of oil used is less than water, so the droplet stays at the bottom of the open well. Finally, the tube which contained the target cells was then moved into a laminar hood, and buffer (Catalog number: SCB-E001M, Qingdao Single-cell Biotechnology, Qingdao, China) was added into the tube for cell lysis.

All Raman spectra were preprocessed by background noise subtraction, baseline correction and normalization to C-H band via LabSpec5 software. The selection of post-SCRS-acquisition cells to be sorted was based on a computer algorithm. Specifically, the sorting was based on C-D/(C-D+C-H); this ratio was calculated via dividing C-D peak area from 2,040 to 2,300 $cm^{-1}$ by the sum of C-D area and C-H peak area from 2,800 to 3,100 $cm^{-1}$. The whole pipeline used here has been made available on GitHub (https://github.com/gongyh/RamanD2O).

***Multiple displacement amplification***

Lysis of the post-FISH-scRACS cells was individually carried out at 65 °C for 15 min with 1.5 μL lysis buffer for each of the one-cell samples, followed by addition of 1.5 μL stop solution to neutralize the lysis buffer. Reaction Buffer and DNA Polymerase were added and the mixture was incubated at 30 °C for 8 hours with 70 °C hot-lid temperature for MDA reactions (Catalog number: SCB-E001M, Qingdao Single-cell Biotechnology, Qingdao, China). Blank control (without any cells) was included to detect and quantify potential contamination. After that, the MDA products were processed for 16S rRNA gene PCR analysis using the 27 F and 1492 R primers (**Table S1**). Once the MDA products of single cell genomic DNA were confirmed to contain single-species 16S-rRNA, they would undergo high-throughput sequencing.

***Library construction and next generation sequencing***

***16S rRNA gene sequencing*** Total genome DNA from seawater sample was extracted using the Fast DNA SPIN extraction kits (MP Biomedicals, Santa Ana, CA, USA) according to the manufacturer's protocols. DNA concentration was measured by UV spectrophotometer Nanodrop NC2000 (Thermo Scientific, Waltham, MA, USA) and its purity was monitored on 0.8% (w/v) agarose gels. 16S rRNA genes of the V3-V4 region were amplified using barcoded 338 F and 806 R (**Table S1**). PCR reactions were carried out in 25 μL reactions with 5 μL of 5× Q5 Reaction Buffer; 5 μL of 5× Q5 High GC Enhancer; 0.5 μL of dNTP Mix (10 mM); 1 μL of each primer (10 μM); 0.25 μL of

Q5 High-Fidelity DNA Polymerase; 2 μL of template DNA and 10.25 μL of $ddH_2O$ (New England Biolabs, USA). Thermal cycling consisted of initial denaturation at 98 °C for 5 min, followed by 30 cycles of denaturation at 98 °C for 10 s, annealing at 50 °C for 30 s, and elongation at 72 °C for 30 s. Finally, 72 °C for 5 min.  Mix same volume of 1 x loading buffer (containing SYB green) with PCR products and operate electrophoresis on 2% agarose gel for detection. PCR products were mixed in equidensity ratios. Then, mixture PCR products was purified with AxyPrep DNA gel extraction kit (Axygen Biosciences, USA). Sequencing libraries were generated using Illumina TruSeq DNA PCR-Free Library Preparation Kit (Illumina, USA) following manufacturer's recommendations and index codes were added. The library quality was assessed on the Qubit@ 2.0 Fluorometer (Thermo Scientific, USA) and Agilent Bioanalyzer 2100 system. At last, the library was sequenced on an Illumina NovaSeq platform and 250 bp paired-end reads were generated.

***Metagenomic sequencing*** Following the manufacturer's instructions, the OMEGA soil DNA kit (Omega Bio-tek, U.S.A) was used to extract the microbial genomic DNA from the seawater samples. A NanoDrop ND-1000 spectrophotometer (Thermo Fisher Scientific, USA) and agarose gel electrophoresis were used, respectively, to evaluate the amount and quality of the extracted DNA. Using the Illumina TruSeq® Nano DNA LT library preparation kit, the extracted DNA from each sample (optical concentration > 2.5 ng/L and total content above 200 ng) was processed to create metagenome shotgun sequencing libraries with 400 bp insert sizes. Each library was sequenced using the Personal Biotechnology Co., Ltd. Illumina NovaSeq platform with the PE150 strategy (Shanghai, China).

***Bacterial one-cell genome sequencing via FISH-scRACS-Seq*** The MDA products from post-FISH-scRACS single-cells were treated with S1 Nuclease (Thermo Fisher Scientific, USA) to degrade the single-stranded nucleic acids, and then purified by Agencourt AMPure XP Beads (Beckman Coulter, USA). A total amount of 0.2 µg DNA per sample was used as input material for the DNA library preparations. Sequencing library was generated using NEB Next® Ultra™ DNA Library Prep Kit for Illumina (NEB, USA) following manufacturer's recommendations and index codes were added to each sample. Briefly, genomic DNA sample was fragmented by sonication to a size of 350 bp. Then DNA fragments were end-polished, A-tailed, and ligated with the full-length adapter for Illumina sequencing, followed by further PCR amplification. After PCR products were purified by AMPure XP system (Beckman Coulter, USA), DNA concentration was measured by Qubit®3.0 Flurometer (Invitrogen, USA), libraries were analyzed for size distribution by Agilent 2100 Bioanalyzer and quantified by real-time PCR (>2 nM). The clustering of the index-coded samples was performed on a cBot Cluster Generation System using Illumina PE Cluster Kit (Illumina, USA) according to the manufacturer's instructions. After cluster generation, the DNA libraries were sequenced on Illumina platform and 150 bp paired-end reads were generated.

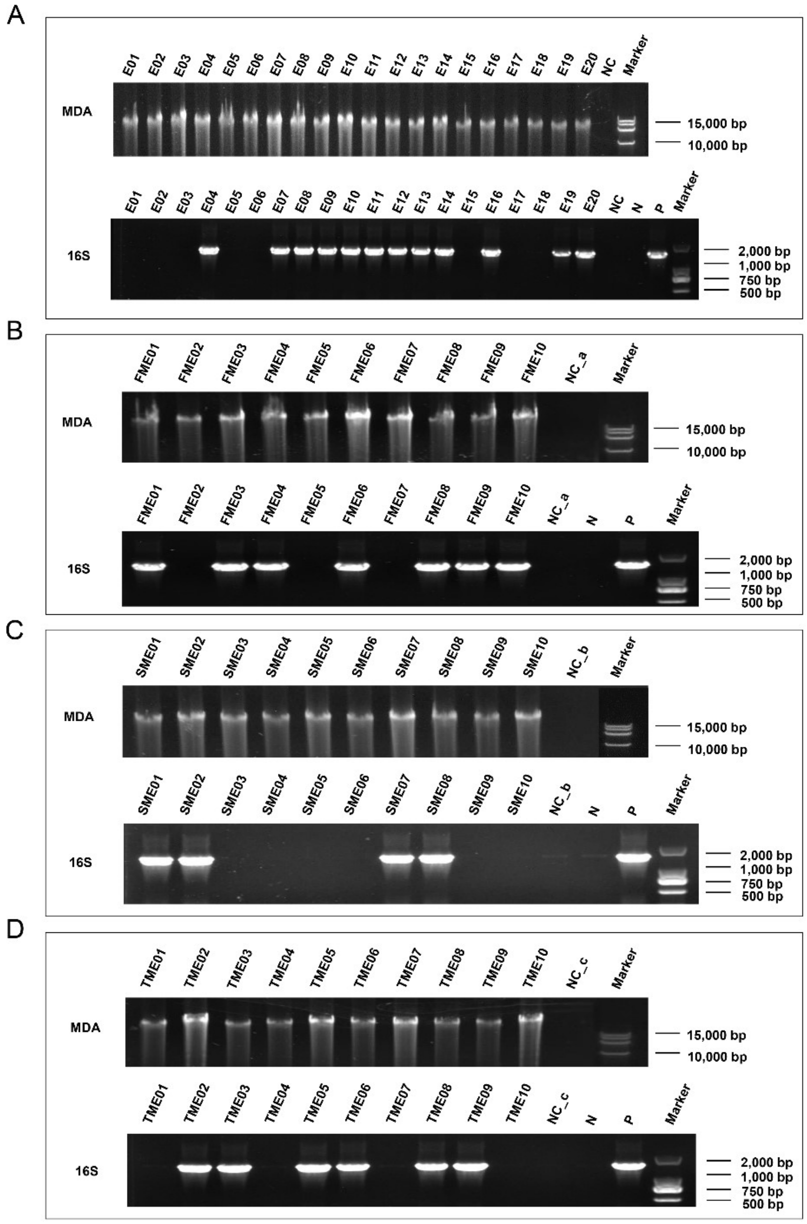


**Figure S1. Agarose gel images of the multiple displacement amplifications (MDAs) and 16S rRNA gene validation processes for target γ-Proteobacteria cells obtained via the FISH-scRACS-Seq.** MDA products and PCR products of the 16S rRNA genes of the target γ-Proteobacteria cells from the pure cultured *Escherichia coli* K-12 DH5α (**A**) and from a four-species mock microbiota (**B-D**, three batches of experiments (B, C and D) were performed). Lane NC, empty droplet (i.e., without cells); Lane N, negative control for PCR (i.e., without adding template); Lane P, positive control for PCR. The non-specific amplification of MDA is due to formation of primer dimers in the MDA reaction.

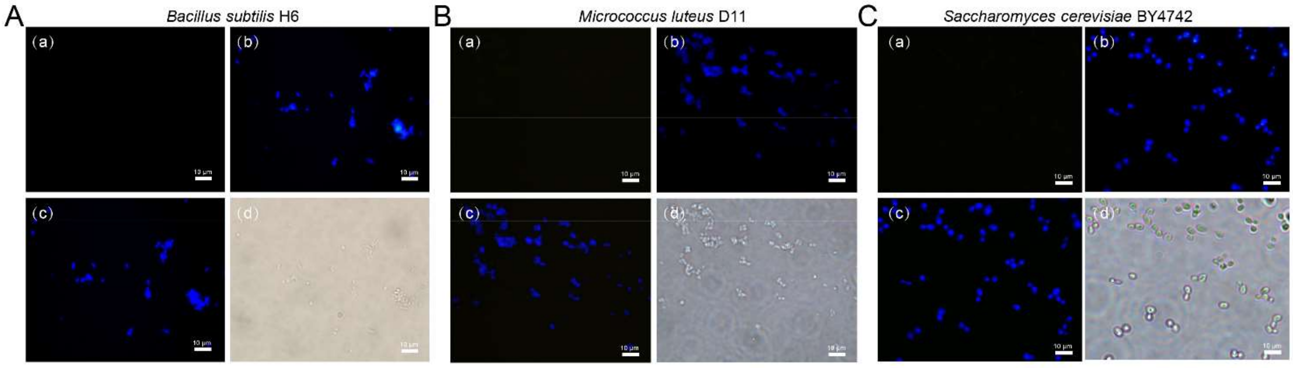


**Figure S2. Validation of the probe GAM42a targeting (A) *Micrococcus luteus* D11 (*Ml, non* γ-Proteobacteria), (B) *Bacillus subtilis* H6 (*Bs, non* γ-Proteobacteria) and (C) one fungus of *Saccharomyces cerevisiae* BY4742 (*Sc, non* γ-Proteobacteria), respectively.** Panel (a) shows the photomicrographs of strain hybridized with HRP-labeled oligonucleotide probes GAM42a; Panel (b) represents DAPI staining (blue) and the same field of view which shows a color combined image recorded by epifluorescence microscopy in Panel (c); Phase-contrast photomicrographs is shown in Panel (d).

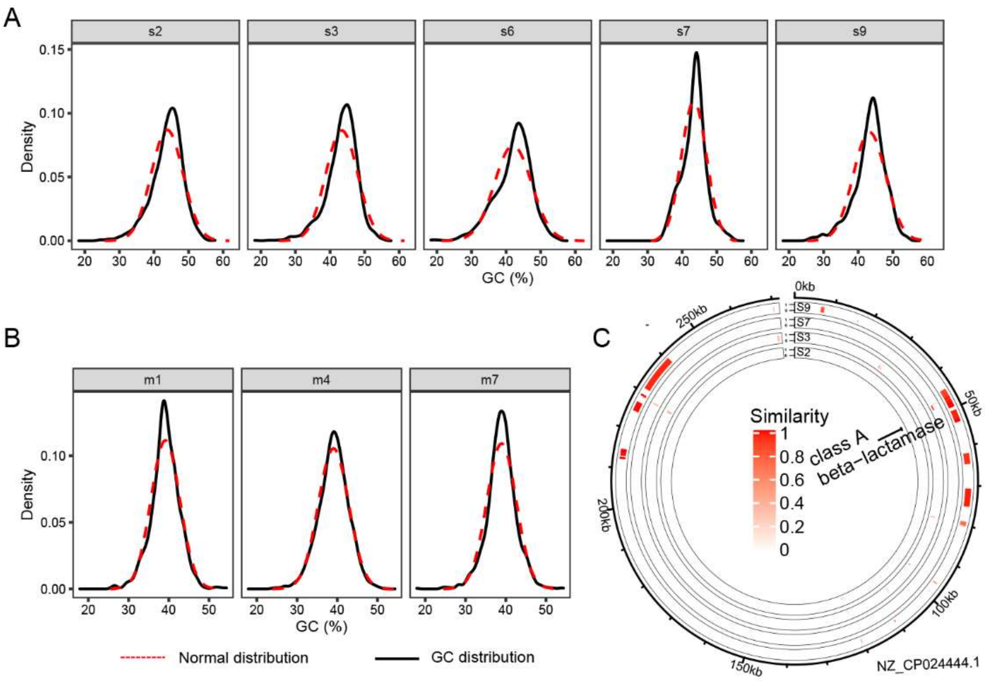


**Figure S3. GC distributions of contigs from the FISH-scRACS-Seq derived SAGs.** GC distributions of the contigs from the SAGs that correspond to FISH-scRACS-sorted cells in soil (**A**) and seawater (**B**), respectively. Black curves represent GC distribution of recovered draft genomes. A sliding window of 200 bp along each contig was used to extract sequence fragments and then calculate GC contents. Red curves show theoretical normal distribution with similar mean and standard deviation to the corresponding GC distribution. The GC contents of these sets of contigs exhibit normal distribution, supporting the integrity of the one-cell assemblies. (**C**) Mapping SAG contigs to a plasmid sequence (accession NZ_CP024444.1) reveals the recovery of a plasmid-encoded antimicrobial resistance gene in the SAG of s9.

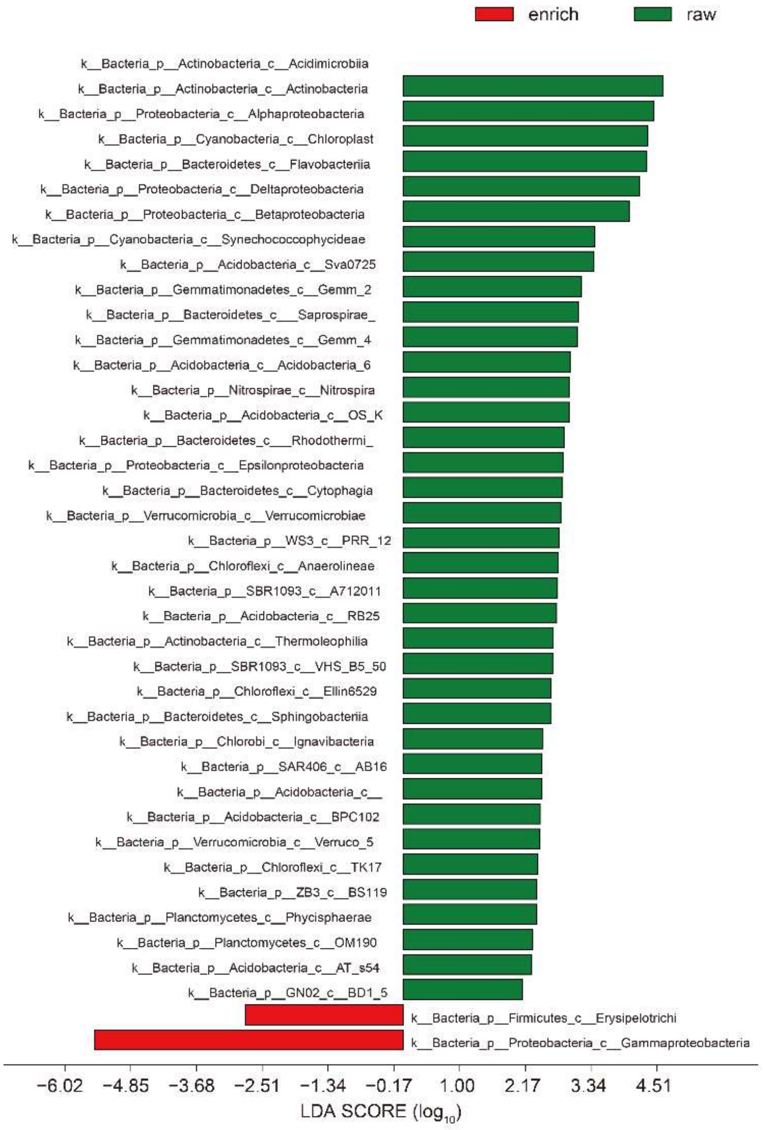


**Figure S4. The bacterial community structures of the original seawater and the $D_2O$-spiked enrichment cultures derived from the Bohai Sea, respectively.**

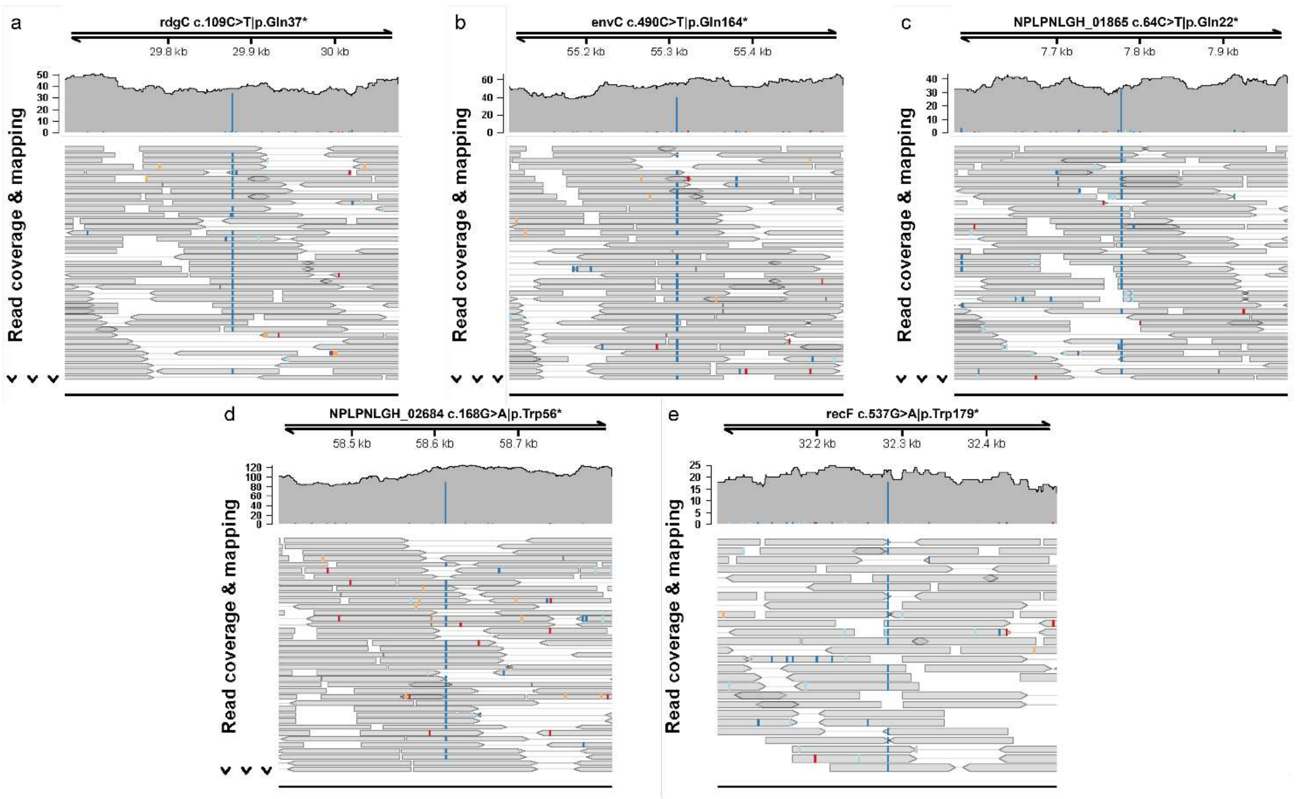


**Figure S5. Read mappings to the MAG1 genome showed sensitive and reliable recovery of chromosomal mutations in SAG1.** Positions of the five SNPs which introduce premature stop codons were shown here. Mismatches in read mappings were highlighted using different colors (not gray).

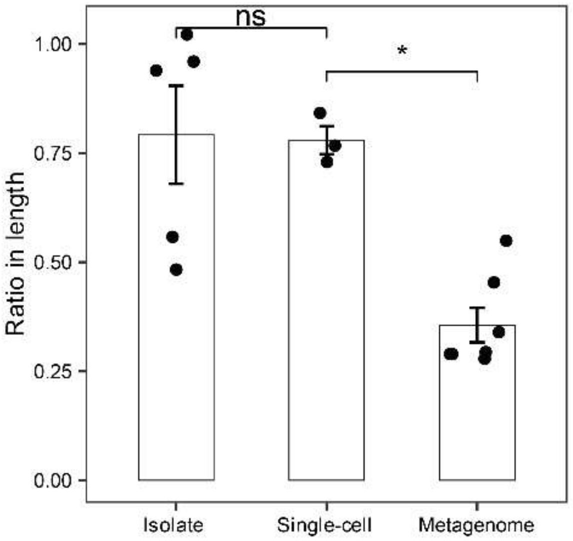


**Figure S6. The insertion sequences (ISs) recovered from bulk sample, SAGs and MAGs, respectively.** Genome sequences of isolated *P. fuliginea* ("Isolate") are retrieved from NCBI RefSeq database. Single-cell genomes ("Single-cell") include SAGs of m1, m4 and m7. MAG genomes ("Metagenome") consist of MAG1-MAG7.

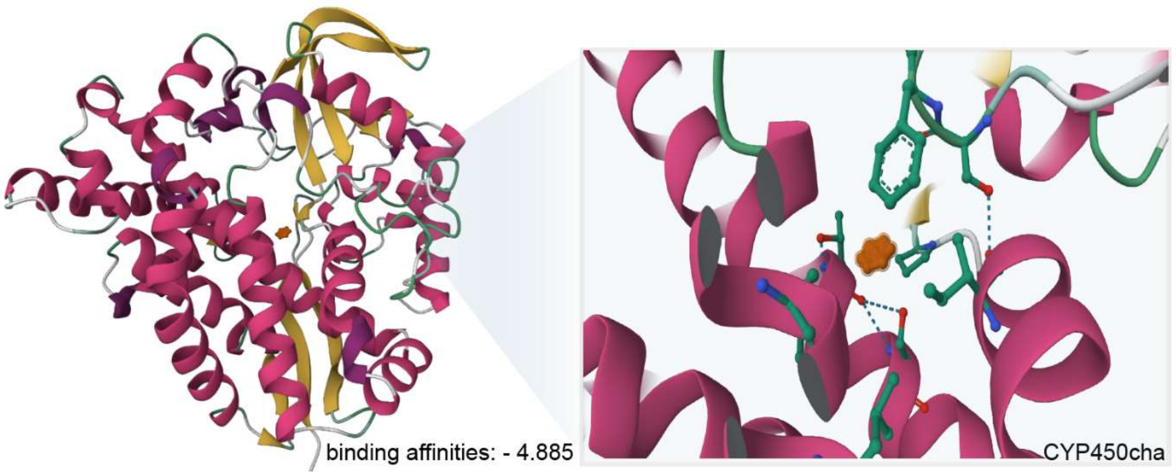


**Figure S7. The protein-ligand docking simulation reveals that cyclohexane (colored in orange) can conjunct with the CYP450cha (AKJ87746.1 from *Acidovorax* spp.).**

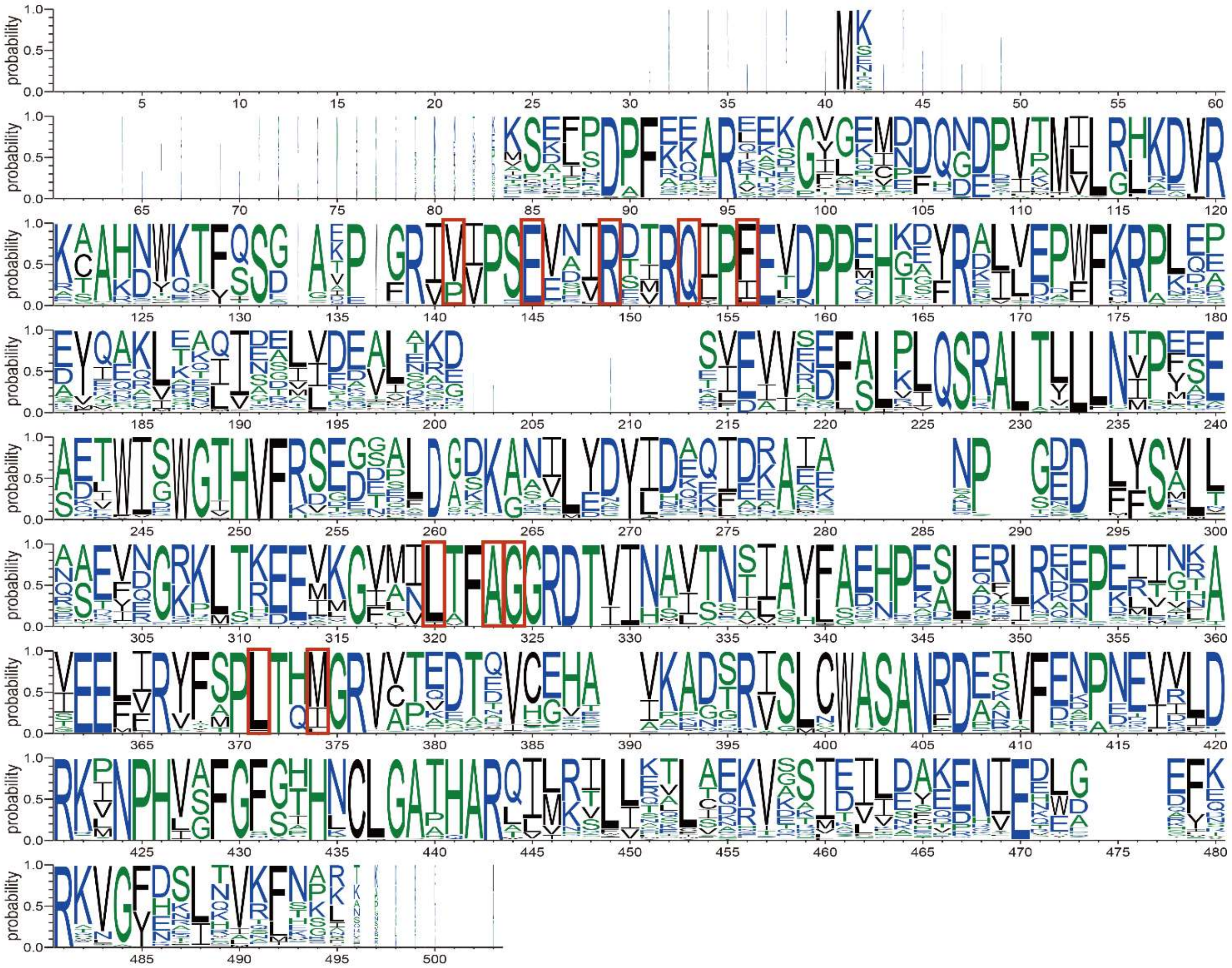

**Figure S8. Sequence logo of CYP236 family proteins selected from the UniRef90 database.** The active sites of G6Me to P450$_{ZoGa}$, which contains all the potential active sites of cyclohexane to P450$_{PsFu}$, are highlighted in red squares.

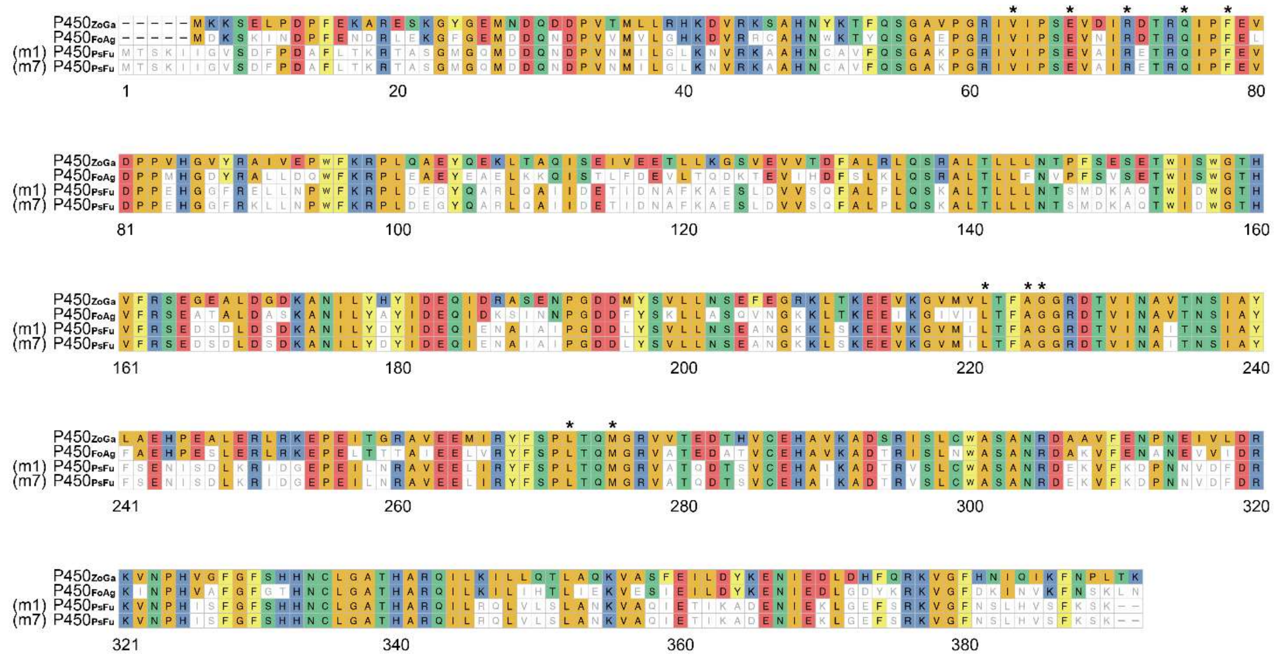


**Figure S9. Multiple sequence alignments of P450$_{PsFu}$ (in SAGs m1 and m7), P450$_{ZoGa}$ and P450$_{FoAg}$.** The active sites of G6Me to P450$_{ZoGa}$ are conserved and marked with black asterisk.

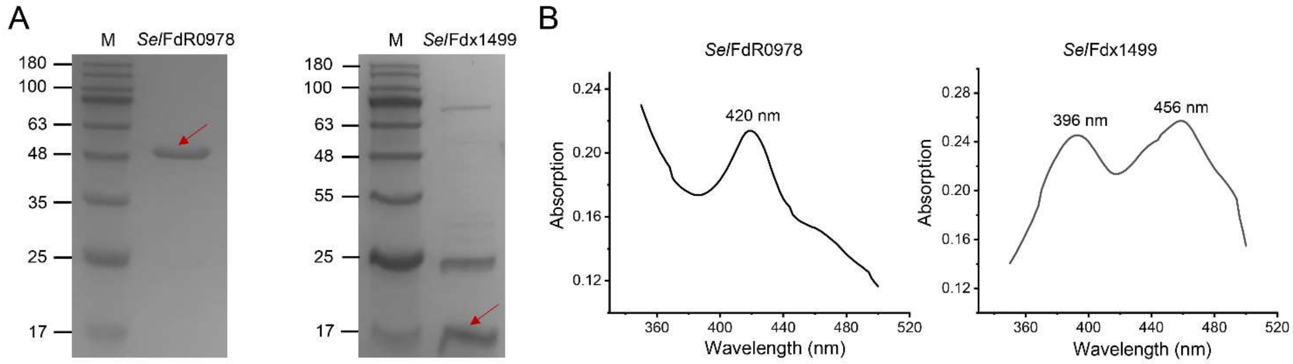


**Figure S10. SDS-PAGE (A) and UV/Vis absorption spectra (B) of *Sel*FdR0978 and *Sel*Fdx1499.**

**Table S1. Probes used in FISH-scRACS-Seq and primers used for PCR amplification of 16S rRNA gene for MDA products.**

| Description | Name | Primer Sequence (5'→ 3') |
|---|---|---|
| **Probe for γ-Proteobacteria** | GAM42a | GCCTTCCCACATCGTTT |
| **Negative probe** | NONEUB | ACTCCTACGGGAGGCAGC |
| **16S rRNA, partial** | 338-F | ACTCCTACGGGAGGCAGCA |
| | 806-R | GGACTACHVGGGTWTCTAAT |
| **16S rRNA, full length** | 27-F | AGAGTTTGATCCTGGCTCAG |
| | 1492-R | GGTTACCTTGTTACGACTT |

**Table S2. Performance of FISH-scRACS-Seq targeting *Escherichia coli* K-12 DH5α (*Ec*).** The GC content and the genome size of reference genome is 50.79% and ~4. 64 Mb (RefSeq: NC_000913.3).

| Assembly | Clean read pairs (million) | Base coverage (%) | Genome fraction (%) | Estimated genome completeness by CheckM (%) | GC (%) | N50 (bp) | Total aligned length (bp) | Largest alignment (bp) | Largest contig (bp) |
|---|---|---|---|---|---|---|---|---|---|
| **E04** | 8.18 | 85.93 | 85.63 | 92.94 | 50.38 | 23,469 | 4,001,748 | 72,054 | 92,992 |
| **E07** | 9.66 | 84.19 | 83.29 | 91.90 | 47.08 | 10,820 | 3,889,692 | 47,576 | 59,782 |
| **E08** | 8.56 | 67.06 | 64.05 | 65.74 | 47.18 | 5,659 | 3,044,418 | 26,381 | 44,542 |
| **E09** | 7.74 | 74.14 | 72.33 | 76.40 | 49.67 | 8,293 | 3,462,974 | 41,688 | 41,688 |
| **E10** | 8.41 | 83.83 | 82.77 | 95.53 | 42.98 | 6,942 | 3,889,423 | 44,616 | 71,901 |
| **E11** | 7.82 | 83.64 | 82.69 | 86.44 | 47.02 | 12,828 | 3,873,805 | 47,338 | 72,858 |
| **E12** | 7.94 | 60.62 | 55.42 | 66.69 | 46.69 | 4,090 | 2,589,749 | 26,980 | 47,355 |
| **E13** | 8.16 | 86.78 | 86.73 | 92.65 | 47.18 | 12,375 | 4,070,110 | 79,710 | 92,539 |
| **E14** | 7.41 | 87.59 | 87.80 | 74.67 | 47.75 | 15,769 | 4,117,474 | 77,932 | 135,496 |
| **E16** | 7.99 | 86.20 | 86.17 | 95.10 | 46.94 | 12,692 | 4,048,088 | 74,094 | 97,151 |

**Table S3. Sequencing and assembly statistics for single-cell genomes from the soil and seawater produced by FISH-scRACS-Seq.**

| FISH-scRACS-sorted samples | | Sequencing | | Assembly | | | |
|---|---|---|---|---|---|---|---|
| | | **Raw read pairs (Million)** | **Clean read pairs (Million)** | **Assembly size (Mbp)** | **Number of contigs** | **N50** | **Number of genes** |
| C-D peak-containing γ-Proteobacteria cells in soil | **s2** | 13.48 | 13.48 | 1.21 | 1,302 | 1,682 | 1,141 |
| | **s3** | 12.73 | 12.73 | 2.21 | 1,370 | 4,662 | 1,928 |
| | **s6** | 11.88 | 11.87 | 3.01 | 930 | 13,288 | 2,931 |
| | **s7** | 11.59 | 11.59 | 2.68 | 91 | 143,031 | 2,404 |
| | **s9** | 13.38 | 13.38 | 2.15 | 863 | 9,450 | 1,925 |
| Cycloalkane-consuming γ-Proteobacteria cells in seawater | **m1** | 10.30 | 10.30 | 4.42 | 852 | 29,235 | 3,853 |
| | **m4** | 11.93 | 11.93 | 2.32 | 2,142 | 5,166 | 1,810 |
| | **m7** | 13.39 | 13.38 | 3.74 | 1,598 | 10,980 | 3,204 |

**Table S4. Carbohydrate-active enzymes in sample m1.**

| Gene in sample m1 | EC# | CAZy class | Signal peptide | Description |
|---|---|---|---|---|
| DBCIMJHN_02445 | 4.2.2.3 | PL7_5 | Y (1-35) | Alginate lyase |
| DBCIMJHN_03826 | 2.4.-.- | GT9 | N | Lipopolysaccharide heptosyltransferase 1 |
| DBCIMJHN_04004 | 3.2.1.80 | CBM38+GH32 | Y (1-26) | Levanase |
| DBCIMJHN_04507 | - | CE12 | Y (1-20) | Rhamnogalacturonan acetylesterase RhgT |
| DBCIMJHN_05125 | 2.4.1.- | GT2 | N | Glycosyl transferase, family 2 |
| DBCIMJHN_05212 | 3.2.1.81 | GH86 | Y (1-22) | Beta-porphyranase A |
| DBCIMJHN_05697 | 4.2.2.- | CBM50+GH23 | Y (1-30) | Membrane-bound lytic murein transglycosylase D |
| DBCIMJHN_06113 | 1.1.99.1 | AA3_2 | N | Oxygen-dependent choline dehydrogenase |
| DBCIMJHN_06202 | 3.2.1.54 | GH13 | Y (1-22) | Neopullulanase |
| DBCIMJHN_06372 | 3.2.1.68 | CBM48+GH13_11 | N | Glycogen operon protein GlgX |
| DBCIMJHN_06416 | 3.2.1.21 | GH3 | Y (1-29) | Beta-glucosidase BoGH3B |

**Table S5. List of CYP236A P450 subfamily enzymes searched from the KMAP metagenomic database.**

| No. | Gene | Protein identity |
|---|---|---|
| 1 | eAquatic_003651446 | 62.2% |
| 2 | eAquatic_002596427 | 61.6% |
| 3 | TARA_GCv2_003731819 | 57.9% |
| 4 | eMarineSediment_001127448 | 58.7% |
| 5 | TARA_GCv2_003214717 | 57.6% |
| 6 | eFreshWater_000936066 | 60.9% |
| 7 | TARA_GCv2_003785908 | 57.9% |
| 8 | TARA_GCv2_004002605 | 59.3% |

**Table S6. List of CYP236A P450 subfamily enzymes searched from the Ocean Microbial Reference Catalog v2.**

| Sample id | Genes | Temperature | Region |
|---|---|---|---|
| TARA_R100000687 | scaffold30530_2_gene25301 | 7.21 | South Pacific Ocean |
| TARA_R110002012 | scaffold57068_1_gene146848;<br>scaffold86101_2_gene214242 | 10.79 | North Atlantic Ocean |
| TARA_R110002051 | scaffold71516_1_gene128954 | 0.22 | Arctic Ocean |
| TARA_R110002049 | scaffold171426_2_gene337933;<br>scaffold192787_2_gene361608;<br>scaffold212459_1_gene383502;<br>scaffold25956_1_gene90675;<br>scaffold7130_1_gene42496 | 8.49 | Arctic Ocean |
| TARA_R110002060 | C2114871_1_gene98357 | 0.10 | Arctic Ocean |
| TARA_R110002074 | scaffold84259_6_gene187075 | 3.11 | Arctic Ocean |
| TARA_R110002072 | scaffold81897_1_gene187239 | 7.53 | Arctic Ocean |
| TARA_R110002096 | scaffold72981_6_gene173149 | -1.30 | Arctic Ocean |
| TARA_R110002124 | scaffold197972_2_gene365002;<br>scaffold9146_5_gene47185 | 1.24 | Arctic Ocean |
| TARA_R110000868 | scaffold1728_8_gene13852;<br>scaffold254352_1_gene510926 | 3.48 | Arctic Ocean |
| TARA_R110002126 | scaffold34643_1_gene107040 | -1.53 | Arctic Ocean |
| TARA_R110000744 | scaffold44106_1_gene98516 | -0.41 | Arctic Ocean |
| TARA_R110000751 | scaffold307927_1_gene436114;<br>scaffold55346_3_gene118730 | 0.99 | Arctic Ocean |
| TARA_R110000737 | scaffold56795_2_gene81598 | -1.33 | Arctic Ocean |
| TARA_R110000765 | C18959726_1_gene609343 | 1.23 | Arctic Ocean |
| TARA_R110002167 | scaffold65492_3_gene185398 | 3.03 | Arctic Ocean |
| TARA_R110002153 | scaffold47696_26_gene134663 | 1.44 | Arctic Ocean |
| TARA_R110001632 | scaffold85373_5_gene187427;<br>scaffold90947_2_gene195217 | 3.20 | Arctic Ocean |
| TARA_R110001599 | scaffold2861_7_gene15501 | 4.20 | Arctic Ocean |
| TARA_R110001606 | scaffold260012_1_gene407756 | 2.50 | Arctic Ocean |
| TARA_R110001583 | C5633139_1_gene407212 | 4.79 | Arctic Ocean |
| TARA_R110001592 | scaffold352891_1_gene651242;<br>scaffold63472_2_gene194345 | 5.29 | Arctic Ocean |

**Supplementary file 1. List of the taxonomy abundance for the seawater samples (A, B and C represent three biological replicates).**

**Supplementary file 2. Genes identified as CAZymes in SAG of m1.**
